# KIF18A in HCC Metastasis: Dual Role in Anoikis Resistance and Chromosome Instability

**DOI:** 10.1101/2024.12.20.629627

**Authors:** Changhao Kan, Weiyu Bai, Xinru Yang, Hayam Hamdy, Xianghui Yang, Xin Liu, Yan Lv, Junling Shen, Jianwei Sun

## Abstract

The study explores the role of KIF18A, a gene linked to whole-genome doubling (WGD) and chromosomal instability (CIN), in the metastatic progression of hepatocellular carcinoma (HCC). We found that KIF18A is a critical link between CIN and anoikis resistance, key factors in HCC metastasis. Through the bulk-seq analysis, we identified the pressures of anoikis and CIN faced by tumor cells during HCC metastasis and identified hub genes central to this process. Our results reveal that E2F activation during HCC progression leads to the transcription of KIF18A, promoting the survival and metastasis of CIN tumors. Overexpression of KIF18A leads to longer tumor life, more micronuclei, and increased survival and metastasis through non-canonical NF-kB activation. Deletion of KIF18A stabilizes the nuclear membrane of the micronucleus, silences cGAS-STING and PI3K-AKT pathways, and inhibits classical NF-kB. The study’s limitations include the need for further animal studies to validate the findings and explore the transient activation of the cGAS-STING pathway. Additionally, future research could focus on the potential therapeutic implications of targeting KIF18A in cancer treatment.

## 1. Introduction

Hepatocellular carcinoma (HCC) is a highly aggressive cancer characterized by abnormal cell differentiation, infiltrative growth, early metastasis, and poor prognosis^1^. Despite significant advancement in liver cancer treatment, the 5-year overall survival rate for HCC patients remains disappointingly low, primarily due to the high rates of metastasis and recurrence^2,3^. These factors contribute to the limited success of current therapeutic approaches and underscore the need for more effective strategies.

The progression of HCC is intricately influenced by anoikis, a biological mechanism that induces programmed cell death to eliminate cells that lose their attachment to the extracellular matrix or neighboring cells (the structure supporting cells)^4,5^. This process is essential for maintaining tissue homeostasis and preventing abnormal cell proliferation. However, dysregulation of anoikis can facilitate cancer metastasis. Cancer cells that evade anoikis can detach from the primary tumor, survive in circulation, and eventually form secondary tumor formation contributing to the spread of the disease ^6,7^.

Chromosomal instability (CIN) is another hallmark of HCC and many other cancers characterized by an abnormal number of chromosomes and increased genetic diversity within cancer cells, which is associated with resistance to therapy and more aggressive tumor behavior^8,9^. The complex interplay between anoikis resistance and CIN in HCC remains poorly understood, but. Emerging evidence suggests that interaction plays a critical role in the metastatic and progressive nature of full understanding of the disease.

This work highlights the significant connection between anoikis resistance and chromosomal instability in hepatocellular cancer, emphasizing its importance. Our study is poised to offer novel insights into the management of HCC, potentially leading to the identification of new therapeutic targets and prognostic biomarkers, ultimately improving the management of HCC and patient outcomes.

## 2. Materials and Methods

### 2.1. Data Collection

The gene expression profile, phenotypic data, and survival data of HCC patients were retrieved from the TCGA-LIHC database via the UCSC Xena database (https://xenabrowser.net/datapages/, accessed September 20, 2022)^10^. To ensure the data integrity, samples with a follow-up period of less than 30 days were excluded from the analysis. Differentially expressed genes (DEGs) were identified using the Deseq2 R package ^11^. The p-values were adjusted using the false discovery rate (FDR) correction method, and the DEG cutoff criteria was set as |log2FC| > 1 and P.adj value < 0.05 (Table. S1). Anoikis-Related Genes (ARGs) were obtained from GeneCards (https://www.genecards.org/, accessed March 16, 2023) ^12^, and genes with a selection criteria of correlation scores >0.4 (Table. S2).

Jvenn (https://jvenn.toulouse.inrae.fr/app/index.html) was used to identify genes associated with anoikis that were significantly expressed in both tumor and normal samples^13^. Univariate Cox regression analysis was performed to identify Anoikis-Related Prognostic Genes (ARPGs), resulting in the selection of a total of 41 ARPGs for subsequent analysis (Table. S3).

We performed PT-CTC and RT-CTC model construction using hepatocellular carcinoma circulating tumor cell single-cell data (CNP0000095)^14^ and primary treatment data from 12 patients with primary HCC and 6 patients with early recurrent HCC (CNP0000650)^15^.

### 2.2. Functional enrichment analysis and Protein-Protein Interaction network of ARPGs

To explore functional roles of ARPGs, Gene Ontology (GO) annotation and Kyoto Encyclopedia of Genes and Genomes (KEGG) pathway enrichment analyses were conducted using the “cluster profiler” package in R^16–18^. The GO terms encompass three components: biological process (BP), cellular component (CC), and molecular function (MF). The results were visualized using the “tidyr” and “ggplot2” R packages with significance set at P.adj < 0.05 and q.value < 0.2. A protein-protein interaction (PPI) network of ARGs was constructed using GeneMANIA (http://www.genemania.org) database^19^.

### 2.3. Clustering of different Anoikis-Related subtypes

Anoikis-related subtypes were identified based on the expression profiles of ARPGs using the R package “ConsensusClusterPlus”. The optimal number of clusters was determined through consensus matrix analysis and cumulative distribution function (CDF) curves, ensuring robust and stable clustering results. Principal component analysis (PCA) was used to visualize and assess the stability of subtypes^20^.

### 2.4. Revealed distinctive hallmark gene sets within the subtypes

Gene set variation analysis (GSVA) was conducted using the R package “GSVA” (Version 1.40.1) ^21^. To calculate the enrichment score for each sample in the gene set, we utilized a pre-defined gene rank. To evaluate the relevant pathways and molecular mechanisms, we downloaded the h.all.v7.4.symbols.gmt subset from the Molecular Signatures Database (MSigDB)^22^. This subset contains curated gene sets representing various pathways and molecular mechanisms. To ensure a comprehensive analysis, we set the minimum and maximum gene set sizes 5 and 5000, respectively. This range allows us to capture gene sets of varying sizes while avoiding excessively small or large sets that could introduce bias.

### 2.5. Composition of the tumor immune microenvironment

The CIBERSORT algorithm was applied to estimate the immune cell composition of the tumor microenvironment (TME) across different subtypes^21,23^. Furthermore, the ESTIMATE algorithm was used to assess the ESTIMATE score, immune score, stromal score, and tumor purity within each subtype^24^.

### 2.6. Construction of ARPGs-related prognostic risk model by LASSO regression

To evaluate the expression patterns of ARPGs, we employed the least absolute shrinkage and selection operator (LASSO) method with a quantization index. This allowed us to identify the most correlated genes, which were then used to construct the ARPGs model using the R package “glmnet”^25^. To optimize the model and reduce overfitting, we applied 10-fold cross-validation.

In this model, a prognostic risk score, referred to as ARS, was computed using the following formula: ARS = ∑(Coef i * exp i) ^26^. The coefficients (*Coef*) represent the weights assigned to each gene based on their contribution to the risk score.

For our study, we divided the LIHC patients into high-and low-risk subgroups based on the median risk score. To assess the performance of the risk score in predicting patient outcomes, we drew a receiver operating characteristic curve (ROC) using the R package “survivalROC”. Additionally, using the “survival” and “survminer” packages in R, we plotted survival curves for the high-and low-risk groups to visualize the differences in survival outcomes.

### 2.7. Construction and verification of a nomogram

A nomogram was constructed using univariate and multivariate Cox regression analyses, incorporating the anoikis risk score (ARS) and clinical data to predict 1-, 3-, and 5-year survival outcomes in HCC. The nomogram was built with the “rms” and “survival” R packages. Calibration curves and decision curve analysis (DCA) were performed to validate the nomogram’s predictive accuracy.

### 2.8. Unsupervised clustering and markers identification

The Seurat^27^ R package was used for unsupervised clustering of PT-CTC and RT-CTC single-cell data. Prior to this, the data had already undergone quality control and normalization. Consequently, we conducted another round of normalization and employed Harmony to mitigate batch effects^28^. To avoid unexpected noise, we only retained protein-coding genes, and genes that were detected in fewer than 10 cells were excluded from downstream analysis. We performed principal component analysis (PCA) on the normalized expression matrix using the highly variable genes identified by the "FindVariableGenes" function. Ultimately, the Seurat "FindAllMarkers" function was employed to identify genes that were expressed in more than 25% of cells in either of the two populations, enabling the detection of cluster-specific expressed genes through pairwise comparisons of clusters.

Identify the genes that show differential expression in each cluster, where the Log2FC value is greater than 1 and the P.adj value is less than 0.05. Select the top 30 genes for annotation using ACT (http://xteam.xbio.top/ACT/index.jsp)^29^, based on the magnitude of Log2FC and the disparity between pct.1 and pct.2.

To cluster all units, choose the principal components (PCs) that account for 90% of the variance.

### 2.9. Copy number variation analysis (CNV)

The inferred CNV result based on the scRNA-seq data was performed by the inferCNV R package (https://github.com/broadinstitute/inferCNV). T cells were used as normal reference cells in PT-CTC and RT-CTC.

### 2.10. Pathway analysis

To confirm the major pathway changes of PT-CTC and RT-CTC during tumor evolution, SCPA^30^ was used to study the Hallmarker gene set, within the pathways showing the largest FC representing the main activated pathways of this cluster.

### 2.11. Cell developmental trajectory

To construct the tumor evolution maps of PT-CTC and RT-CTC, we used the slingshot^31^ package for pseudotime analysis. In order to reduce the noise impact of complex environments, we use k-means to reclassify tracks that are consistent with the original tracks. To ensure alignments, NB-GAM was used in the tradeSeq^32^.

### 2.12. Cell-cell interaction analysis

To analyze cell-cell interactions between different cell types, we used cellchat^33^ to identify the significantly ligand-receptor pairs within PT-CTC and RT-CTC HCC cells.

### 2.13. Cell line experiments

HCC cell lines (Huh7 and HepG2) and the 293T cell line were purchased from the American Type Culture Collection (ATCC, Manassas, VA, USA). All cell lines were cultured in Dulbecco’s modified Eagle’s medium (DMEM) containing 10% fetal bovine serum (FBS) and levofloxacin. The cells were incubated at 37 °C in a humidified atmosphere containing 5% CO2. Monthly testing for Mycoplasma.

For lentiviral transduction, HepG2 and Huh7 cells were cultured in six-well plates to 70% confluence and infected with lentivirus, which was prepared using peg8000 (Cat#60304ES76, Yeasen).

### 2.14. Configuration of ultra-low adhesion plates and acquisition of anoikis-resistant cell lines

We prepared a polyhema (Cat#ST1582, Beyotime) solution with anhydrous ethanol at a concentration of 120 mg/ml, following the methodology of S-S Liau et al^34^. We diluted it 1:10 with 95% ethanol and added 1.5 ml of the dilution to each well of a 6-well plate, which was subsequently heated and dried for 6 hours to produce ultra-low-adhesion plates.

The plates were rinsed twice with PBS and once with DMEM prior to use. 5x10^6 HepG2 cells were introduced into each well, and HepG2 AR cells were perpetually grown for 7 generations over a span of 14 days to yield HepG2 AR cells. Subsequently, HepG2 AR cells were placed in normal adherent dishes and cultivated for 7 generations over a period of 14 days to obtain stable HepG2 AR adherent cells.

### 2.15. Expression vector and transfection

The KIF18A-coding sequence was PCR amplified from Cloning vector-KIF18A (Cat#SP-103968, BRICS) and using gene-specific primers modified to include the appropriate cognate arm sites at their 5′ end. The purified PCR product was ligated by homologous recombination (Cat#10923ES50, Yeasen) with the Phage-3xFlag-egfp-puro vector or Phage-3xFlag-puro vector digested by BamHⅠ and XhoⅠ.

[UTBL]

### 2.16. shRNA-mediated KIF18A knockdown

Lentiviruses encoding non-silencing shRNA (vector-shRNA, control) and KIF18A-specific shRNA (knockdown) were generated. The shRNA targeting sequences used were as follows:

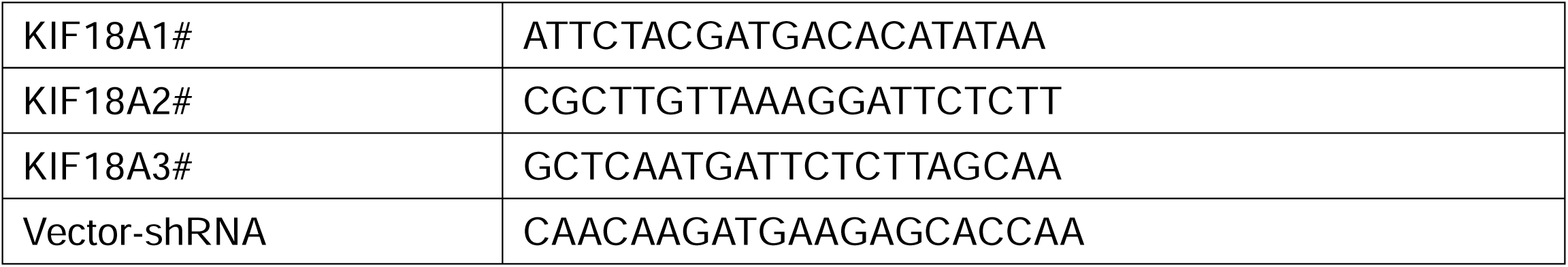

### 2.17. Quantitative real-time PCR and Western blot

Total RNA was isolated, and the cDNA was prepared, amplified, and detected in the presence of SYBR as previously^35^.

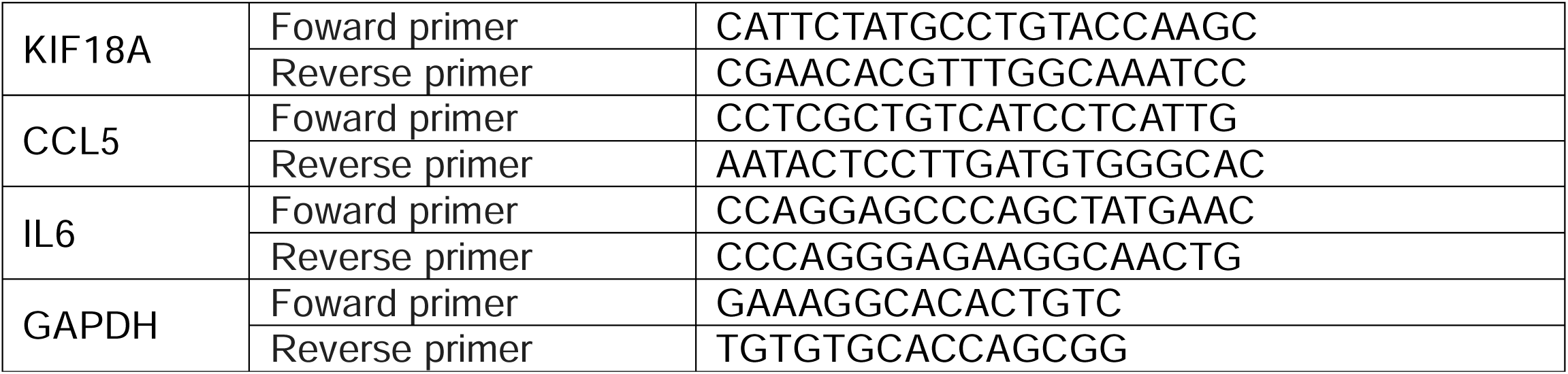

Cell lysates and supernatants were resolved by electrophoresis, transferred to a polyvinylidene fluoride membrane, and probed with antibodies.

| Antibodies | Source | Identifier | Diluted |
| --- | --- | --- | --- |
| KIF18A | Santa Cruz | sc-390600 | 1:500 |
| p-STING(Ser366) | Affinity | AF7416 | 1:500 |
| NF-kB(P65) | CST | #3034 | 1:1000 |
| p-NF-kB(P65)(Ser536) | CST | #3031S | 1:1000 |
| NF-kB(P52) | Santa Cruz | sc-7386 | 1:500 |
| STAT3 | Selleck | F0200 | 1:1000 |
| p-STAT3(Tyr705) | Selleck | F0157 | 1:1000 |
| STAT1 | ZenBio | R25799 | 1:1000 |
| p-STAT1(S727) | HuaBio | ET1611-20 | 1:1000 |
| TBK1 | Abclonal | A3458 | 1:1000 |
| p-TBK1(Ser172) | Selleck | F0255 | 1:1000 |
| BCL-2 | MCE | HY-P80566 | 1:1000 |

### 2.18. Data processing

All the analysis methods and R packages were implemented by R version 4.3.3. The t-test or Mann–Whitney U test was used to compare the data of two groups. All differences among and between groups were considered statistically significant at p-values ofL<L0.05 (*, pL<L0.05; **, pL<L0.01; ***, pL<L0.001).

## 3. Results

### Identification, Functional Analysis, and Correlation with Prognosis of Anoikis Resistance and CIN in Liver Cancer

To identify ARGs with differential expression overlap, we initially identified DEGs (|log2FC| > 1.0, p.adj < 0.05) in TCGA-LIHC and obtained 485 ARGs from the GeneCards database. We found 126 ARGs with differential expression, as shown in the Venn diagram (Fig. S1A). As indicated in Fig. 1B, univariate Cox proportional hazards regression analysis revealed that 41 ARGs were significantly associated with prognostic outcomes. Some of these are highlighted in the volcano plot (Fig. 1A). To explore the biological significance of these ARPGs, we performed GO analysis and KEGG pathway enrichment (Fig. 1C). The results indicated that most of the genes were involved in organelle fission, nuclear division, and mitotic nuclear division in the BP category. In the CC category, the primary locations for these genes were the chromosomal region, chromosome, centromeric region, and midbody. In terms of MF, the genes were primarily involved in binding tubulin to stabilize microtubules interacting with cell adhesion molecules to protect cells. KEGG pathway analysis revealed that ARPGs were primarily involved in the PI3K-Akt signaling pathway, cancer microRNAs, and focal adhesion, which is important for cell-cell connection.

**Figure 1:**
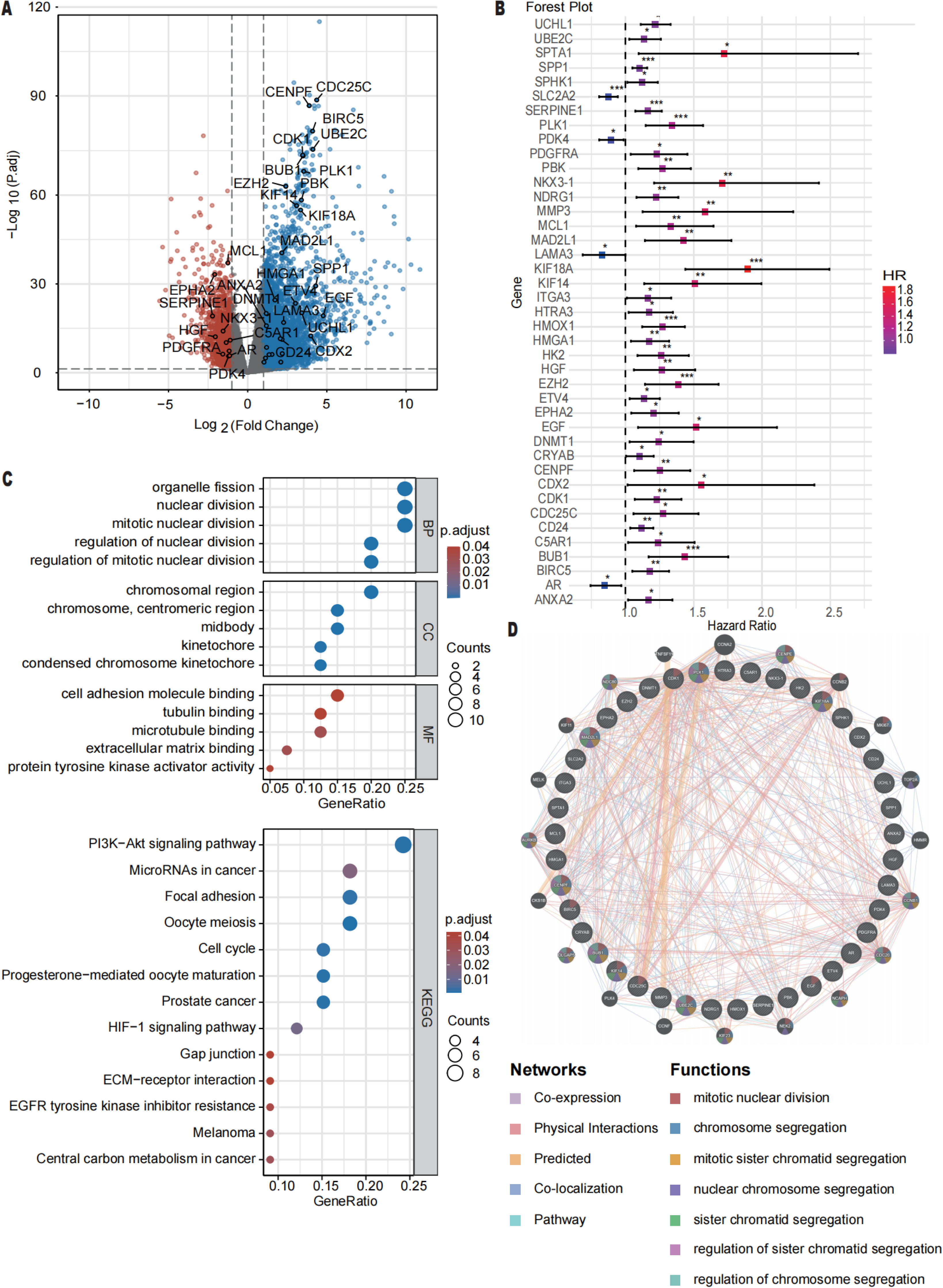
Analysis of ARGs in hepatocellular carcinoma (HCC) (A) The volcano plot depicts differentially expressed genes (DEGs) from the TCGA-LIHC dataset, highlighting those with significant expression changes (|log2FC| > 1.0, adjusted p-value < 0.05). (B) Univariate Cox proportional hazards regression analysis identifies 41 ARGs that are significantly associated with survival outcomes in HCC. (C) Gene Ontology (GO) and Kyoto Encyclopedia of Genes and Genomes (KEGG) pathway enrichment analyses for the ARGs. (D) The protein-protein interaction (PPI) network, constructed using the GeneMANIA database.

We employed the GeneMANIA database to construct a protein-protein interaction network for ARGs (Fig. 1D) and found that these genes significantly contribute to processes such as chromosome segregation, sister chromatid segregation, nuclear chromosome segregation, and regulation of sister chromatid segregation. These findings provide critical insights into the physiological roles of ARPGs in cellular processes related to cell division and genomic stability. Furthermore, to enhance the clinical relevance of these ARPGs, we performed unsupervised clustering analysis on the TCGA-LIHC dataset (OS. Time ≥ 30 days) and investigated their biological processes and immune cell infiltration (Fig. S1B-S1F and S2). This analysis also raises an important question for our research: why are ARPGs closely related to chromosome segregation and genomic stability processes? Addressing this question may provide valuable insights into the mechanisms underlying liver cancer progression.

### Liver Cancer Prognosis Risk Assessment Based on the ARS Model

We randomly selected 241 samples from the LIHC dataset for the training set and 100 samples for the test set. Subsequently, we performed LASSO regression analysis using ARPGs, which led to the identification of six genes. The risk score was calculated using the following formula: ARS = (0.173030087 * expression level of KIF18A) + (0.063695246 * expression level of SPP1) + (0.014878703 * expression level of PLK1) -(0.037960223 * expression level of SLC2A2) + (0.001804856 * expression level of EZH2) + (0.121272810 * expression level of MMP3). A significant link association was found between these six identified genes (KIF18A, SPP1, PLK1, SLC2A2, EZH2, and MMP3) and the survival outcomes of individuals with LIHC, suggesting their potential as prognostic markers (Fig. S3A). Furthermore, Fig. S3B illustrates the relationships between patient status, ARPGs clusters, and ARS. Fig. S3C demonstrates that LIHC patients with the ARPGs-C1 subtype exhibit an elevated ARS. This means that the ARPGs-C1 subtype may be a sign of a negative prognosis.

Based on the median ARS, we classified HCC patients into low-risk and high-risk groups. In both the training and test datasets, patients with a high ARS had a significantly shorter OS compared to those with a low ARS. Time-dependent ROC curves were generated for both datasets. These curves demonstrated the ability of our ARS model to predict higher mortality rates in LIHC patients with increasing ARS levels in both the train and test datasets. The curves shown in Fig. 2A–C confirm that the ARS model accurately predicts higher mortality as ARS levels increase in both datasets. Additionally, we conducted a comprehensive analysis of the data from TCGA-LIHC and GSE14520, which yielded consistent results (Fig. 2D).

**Figure 2:**
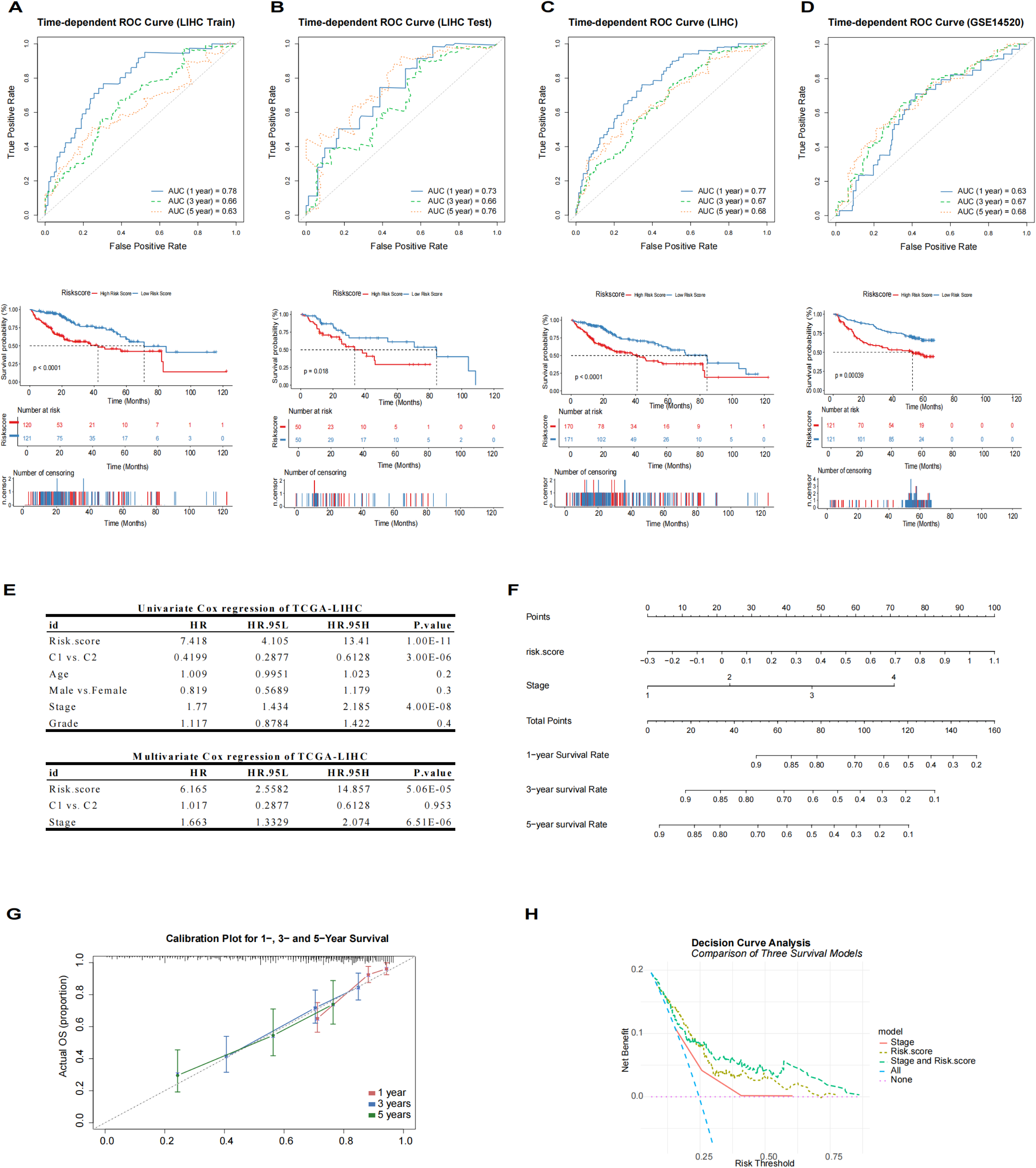
Assessment of the ARS Model’s Predictive Power and Prognostic Factors in LIHC. (A-D) Time-dependent ROC curves for the LIHC train, test, LIHC, and GSE14520 datasets at 1, 3, and 5 years. (E) Univariate and multivariate Cox regression analyses identify the risk score, subtype, and tumor stage as significant prognostic factors in LIHC. (F) Nomogram integrating the ARS model with clinicopathological parameters. (G) Calibration plots for the nomogram at 1, 3, and 5 years, validating the model’s predictive accuracy. (H) DCA detects nomogram performance comparing the predictive accuracy of the nomogram with a single predictor model.

Specifically, univariate Cox regression analysis revealed that the risk score, subtype, and stage were significant risk variables for patients with LIHC. The multivariate Cox analysis further demonstrated that both the ARS model risk score and tumor stage can independently behave as adverse prognostic factors (Fig. 2E).

We developed a nomogram (Fig. 2F) to integrate the ARS model with clinicopathological parameters, providing a tool for clinicians to estimate the survival time of LIHC patients. The calibration curves for the nomogram at 1, 3, and 5 years showed the model’s exceptional predictive efficacy (Fig. 2G). Additionally, a Discriminant Correlation Analysis (DCA) was performed to compare the prediction accuracy of the nomogram with a model that uses only a single predictor (Fig. 2H). The DCA results shown in Fig. S3D confirm that the nomogram outperforms single predictors in predicting OS at 1, 3, and 5 years.

Although extensive research on ARPGs, six specific genes were identified: KIF18A, SPP1, PLK1, SLC2A2, EZH2, and MMP3, which play crucial roles in LIHC prognosis. Notably, KIF18A was strongly associated with the predictive risk score and chromosomal instability in cancers; it is thought to help tumor cells resist anoikis and maintain chromosomal instability, which is essential for tumor progression^36^. Further investigation into the precise mechanisms by which KIF18A contributes to tumor cell resistance and chromosomal instability.

### Single-Cell Data Analysis Reveals the Metastatic Niche in Liver Cancer

We analyzed single-cell data from circulating tumor cells (CTCs) in HCC (CNP0000095) and data from 12 patients with primary HCC and 6 patients with early recurrent HCC undergoing primary treatment (CNP0000650). This analysis aimed to uncover the relationship between ARPGs, tumor metastasis, and chromosomal instability. Tumors and adjacent non-tumor tissues from primary HCC were categorized separately. Tumors from primary HCC and their paired adjacent non-tumor tissues were designated as PT and primary non-tumor (PNT), while those from early recurrent HCC were labeled as RT and recurrent non-tumor (RNT).

To construct a comprehensive atlas of tumor metastatic niches, we utilized Seurat for cell classification and gene identification. The Uniform Manifold Approximation and Projection (UMAP) method was applied to visualize PT-CTC and RT-CTC (Fig. 3A, 3B, and S4A). For all cells except CTC like cell-Primary containing CTC, immune cells highly expressed CD45 (PTPRC), while non-immune cells were EPCAM (Fig. 3A and 3B). Since CTC cells can form heterotypic clusters with immune cells and uptake CD45 from extracellular vesicles^37,38^. CTC like cell in Primary group showed partial admixture with cells from adjacent liver in both the PT-CTC and RT-CTC models (Fig. S4A). Overrepresentation of these cells in certain patients challenges the hypothesis that metastatic potential resides in a small cell cluster ^39^ (Fig. S4B).

**Figure 3:**
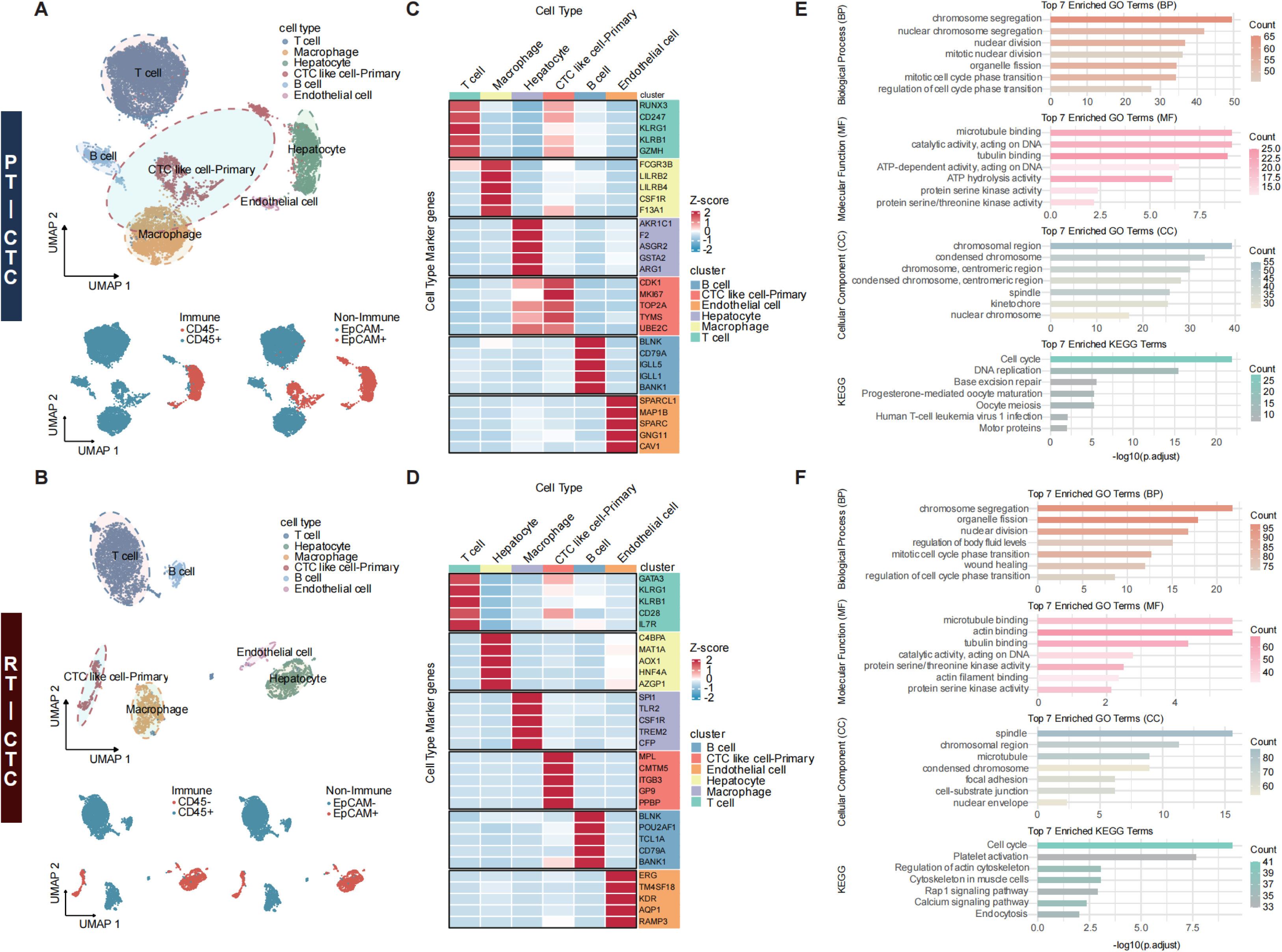
Cellular Composition and Molecular Signatures in Primary and Recurrent HCC. (A-B) UMAP analysis of immune and non-immune cells in both PT and RT HCC. (C-D) Marker gene expression across various cell clusters highlights the cellular composition within PT-CTC and RT-CTC tumor samples. (E-F) GO and KEGG enrichment analyses for CTC like cell-primary.

CTC like cells showed some degree of T cell marker gene expression (Fig. 3C, 3D, and S4C), suggesting a degree of immune cell mixing. inferCNV, GO, and KEGG further supported this conclusion. Specific copy number variants (CNV) were not significant in CTC like cell-Primary clusters (Fig. S4D), but CTC like cell-Primary was enriched in DNA, chromosomes, and cell cycle compared to other clusters (Fig. 3E and 3F).

### Identification of Genuine CTCs and Their Association with Hepatocellular Carcinoma Cells

To explore the relationship between true CTC like cells and malignant hepatocellular carcinoma cells, non-immune (EPCAM+) cells were recharacterized. Immune cells from the first clustering were identified as CTC like cells in both PT-CTC and RT-CTC (Fig. 4A, 4B, and 4C). consistent with prior observation, CTC and CTC like cells can partially mimic immune cells. In the PT-CTC model, some CTC like cells were clustered together with hepatocytes, forming a unique population. However, in the RT-CTC model, only a small subset of CTCs clustered with Hepatocyte-RAB17. This may reflect the limited enrichment of CTC from patients with BCLC stages of A and B, who had not undergone extensive Darwinian selection in recurrent tumor ecology. In the PT-CTC model, CTC like cells up-regulated fibroblast-related genes, while CTCs and Hepatocyte-RAB17 in the RT-CTC model up-regulated hepatocyte markers, potentially reflecting the pre-metastatic EMT and the post-metastatic MET cascade^37^ (Fig. 4D).

**Figure 4:**
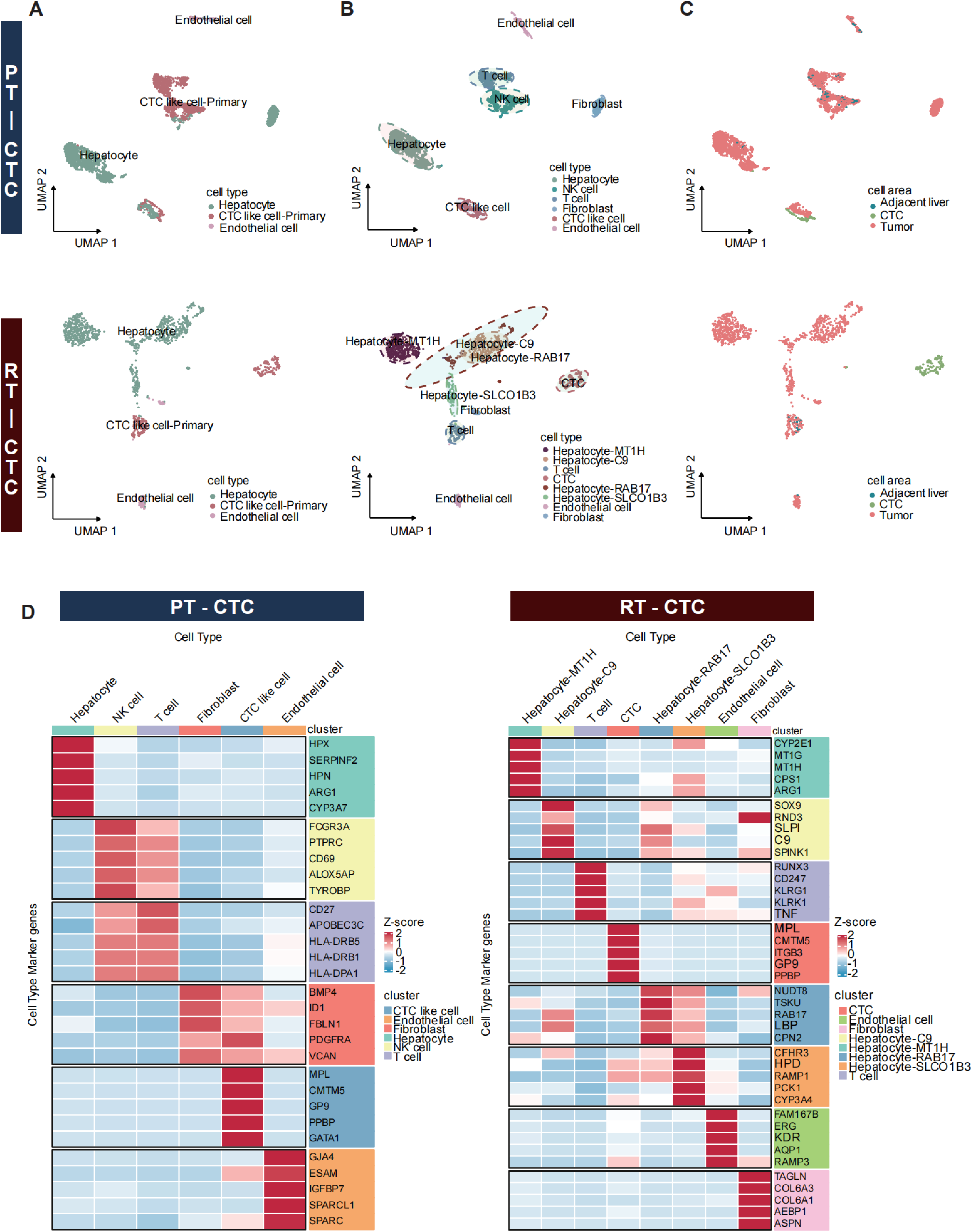
Characterization of CTC-like Cells in Hepatocellular Carcinoma. (A) UMAP plots of PT-CTC and RT-CTC hepatocellular carcinoma models generated from the initial clustering of EPCAM-positive cells. (B) UMAP plots of PT-CTC and RT-CTC hepatocellular carcinoma models generated from the initial clustering of EPCAM-positive cells, with a focus on different cellular components. (C) UMAP plots of PT-CTC and RT-CTC hepatocellular carcinoma models generated from the initial clustering of EPCAM-positive cells, with an emphasis on cellular area. (D) Re-clustering analysis of cell marker genes of the gene expression characteristics in different cell populations.

### Mapping of Tumor Progression Trajectories

CTC like cells and liver cells identified in PT-CTC and RT-CTC were reclassified (Fig. S5A and S5B). Using SCPA, we analyzed their significant pathways (Fig. 5A and Table. S4). and used Slingshot for trajectory tracking (Fig. 5B). In the PT-CTC model, we use the CTC cell as the end point of differentiation, whereas in the RT-CTC model, the CTC cell serves as the starting point. For the PT-CTC model, we have done two tracks, one for the ultimate differentiation of liver cells that are primarily activated by MYC Target V1 and the other for the Heme Metabolism activation of the CTC. Researchers have found that MYC Target V1 activation drives growth^40^, whereas the Heme Metabolism aids the tumor in overcoming anoikis challenges during transplantation and coping with the stress of blood flow shear pressures ^41^. According to the RT-CTC model, we identified a single trajectory, supporting the hypothesis that recurrent tumors often originate from early primary cancer dessemination^42^.K-means clustering confirmed consistency with original trajectories (Fig. S5C). The NB-GAM analysis in tradeSeq identified six genes with the highest trajectory relevance (Fig. 5D).

**Figure 5:**
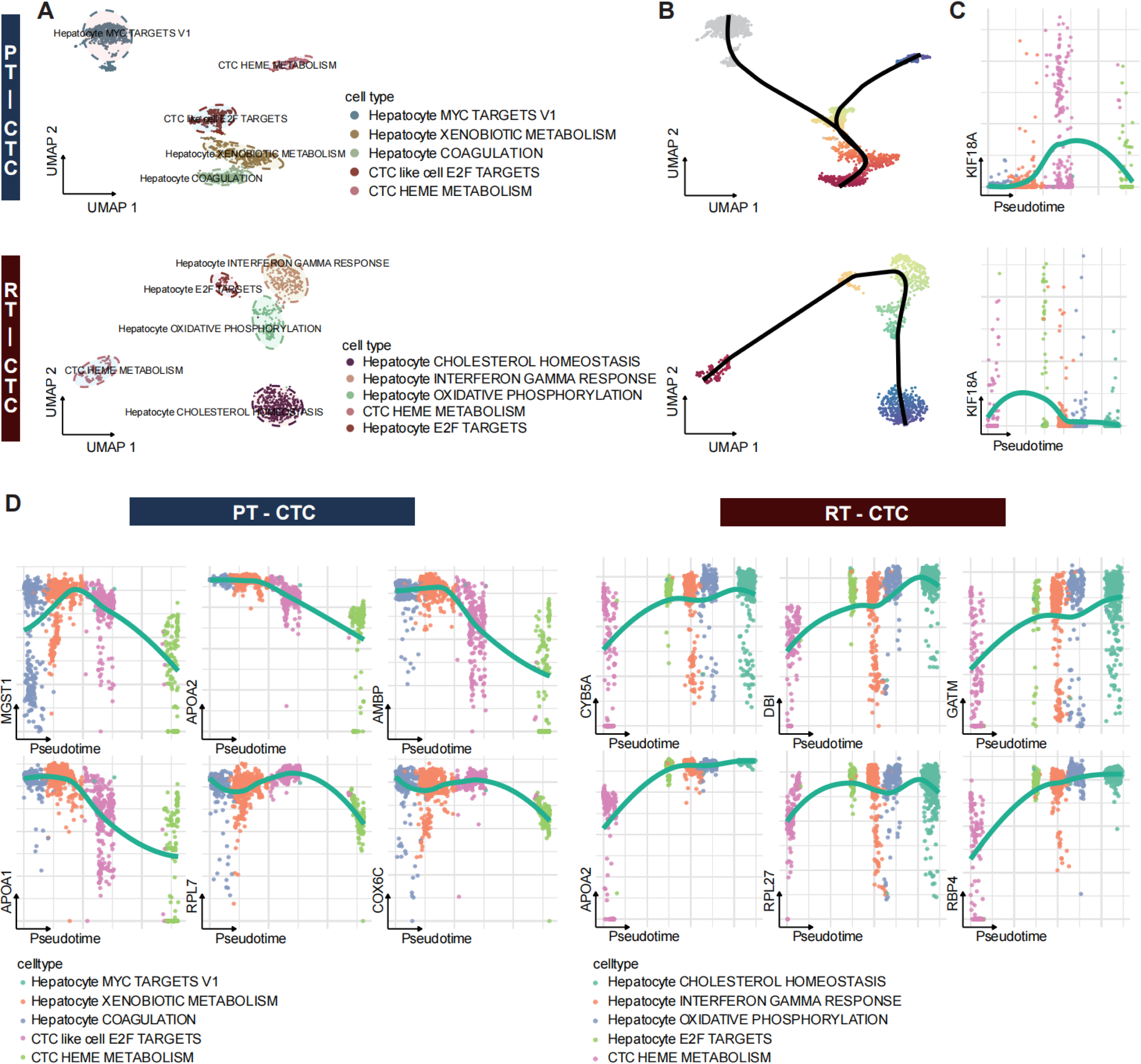
Trajectory Analysis and Gene Expression in CTC-Like and Hepatocyte Cells. (A) Significant pathway analysis for the significant pathways in CTC-like and hepatocyte cells. (B) Trajectory tracking with Slingshot in the PT-CTC and RT-CTC models from hepatocytes to CTC cells. (C) The expression trajectory of KIF18A highlights in both PT-CTC and RT-CTC models. (D) Top gene expression from NB-GAM from the tradeSeq package identifies and displays the six genes with the highest trajectory relevance.

Interestingly, we noticed that the gene KIF18A, which we’re examining in both the PT-CTC and the RT-CTC models, experiences an upgrade in the evolution trajectories of both CTC and CTC like cells (Hepatocyte E2F TARGETS in RT-CTC) (Fig. 5C), and its activation patterns align more closely with the transcription pattern of proliferation-sleep-settlement during tumor transplantation^39^. Combined with previous reports, we have recognized the role of E2F target genes in the process of wire splitting. Activation of E2F promotes KIF18A transcription^43,44^, and in addition, these genes can cause retardation, chromosome rupture, and absence, leading to CIN^45,46^. Additionally, KIF18A is a necessary condition for the survival of WGD+/CIN cells^36,47^. We posit that KIF18A significantly influences the tumor transfer process, facilitating tumor dissemination by alleviating the mounting pressure of increasing chromosomal instability during tumor transplantation.

### The Role of CCL5 in CTC-Like Cell Analysis and Its Impact on Tumor Metastasis Signaling Pathways

We conducted cellchat analysis for PT-CTC and RT-CTC models (Fig. 6A) and visualized pathway activity (Fig. S6A). The chemokine CCL5 is a critical mediator of immune evasion in CTC of patients with hepatocellular carcinoma^14^. Therefore, we focused on the activation of the CCL pathway in PT-CTC and RT-CTC, where CTC actively participated in most activities within the CCL signaling pathway, both as a ligand and a receptor (Fig. 6B and S6B).

**Figure 6:**
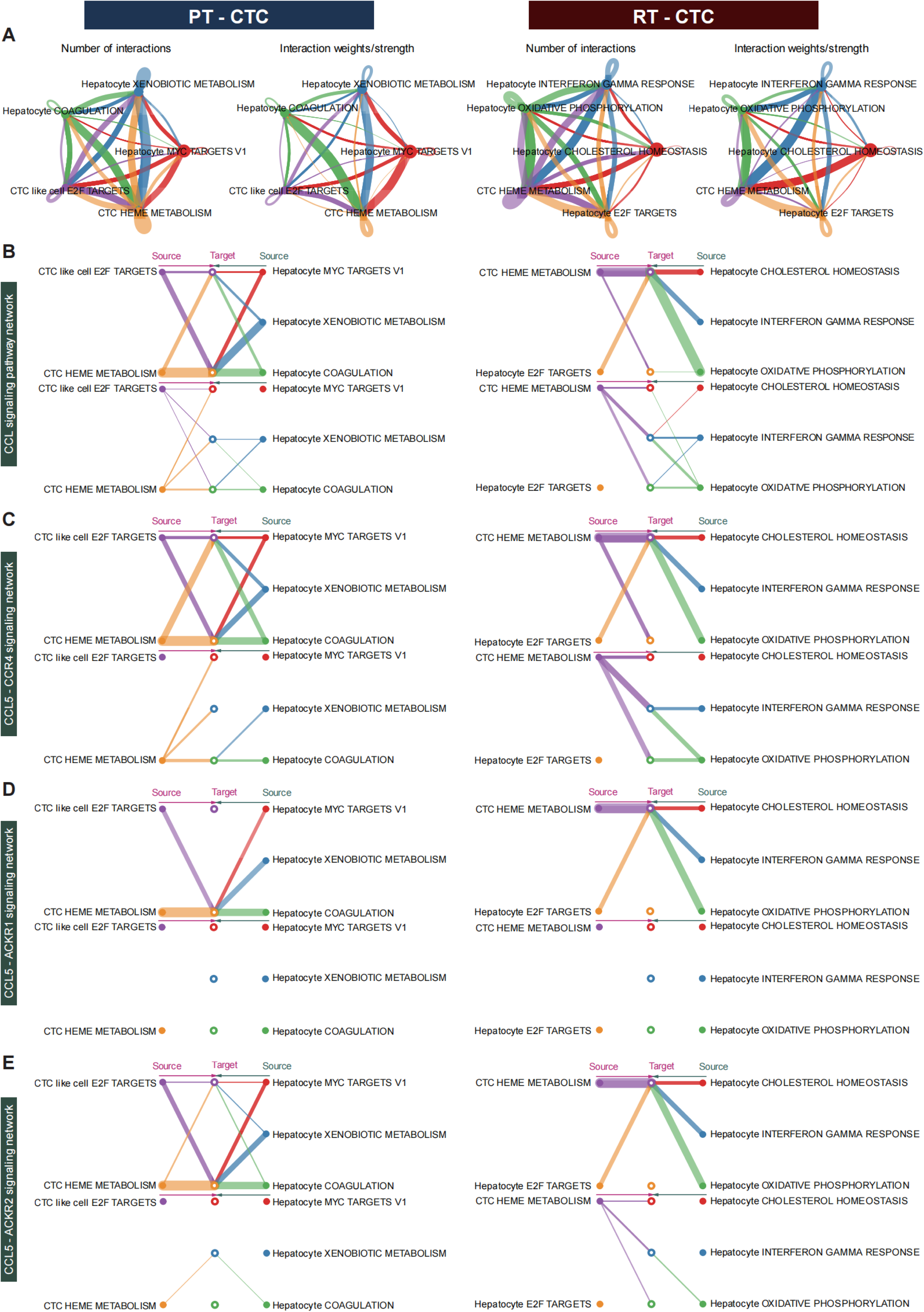
Analysis of CCL Signaling Pathway in PT-CTC and RT-CTC Models. (A) Cell Chat analysis of PT-CTC and RT-CTC for the different cell types within the PT-CTC and RT-CTC models. (B) CCL signaling pathway activation of the CCL signaling pathway in both PT-CTC and RT-CTC. (C) CCL5-CCR4 signaling network between CCL5 and its receptor CCR4. (D) CCL5-ACKR1 signaling network between CCL5 and its receptor ACKR1. (E) CCL5-ACKR2 signaling network between CCL5 and its receptor ACKR2.

To understand the mechanism underlying the activation of the CCL signaling pathway, we examined ligand-receptor interactions, which included CCL5-CCR4, CCL5-ACKR1, and CCL5-ACKR2, as depicted in Figures 6C, 6D, and 6F. Upon analyzing ligand-receptor activation in PT-CTC and RT-CTC, we identified robust interactions between CCL5 and CCR4 involving both CTCs like cells and CTC. Conversely, in RT-CTC, the engagement of hepatocyte E2F TARGETS as receptors showed a relatively weaker intensity (Fig. 6D). According to previous studies, the activation of CCL5-CCR4 promotes EMT transition, facilitating metastasis^14,48^. In PT-CTC, CTC like cells undergo a more pronounced EMT, whereas in RT-CTC, they exhibit MET. This highlight underscores the diverse cellular responses in the context of CCL5-CCR4 activation.

### The Role of KIF18A in Hepatocellular Carcinoma Metastasis and Its Impact on CTC Behavior

To investigate the potential role of KIF18A in HCC metastasis, we initially selected appropriate cell lines based on basal expression levels of KIF18A from the HPA database^49^: HepG2, with lower KIF18A expression, and Huh7, with higher expression (Fig. 7A). Transcription and protein level assessments confirmed differential expression of KIF18A in HepG2 and Huh7 cells (Fig. 7B). For experimental validation, we constructed HepG2 and Huh7 cell lines with KIF18A overexpression (OE) and shRNA-mediated focusing, focusing on HepG2 KIF18A OE and Huh7 KIF18A sh3# cell lines (Fig. 7C, 7D, S7A, and S7B). Our finding revealed that Huh7 KIF18A sh3# cells exhibited increased susceptibility to apoptosis, corroborated via flow cytometry (Fig. 7E).

**Figure 7:**
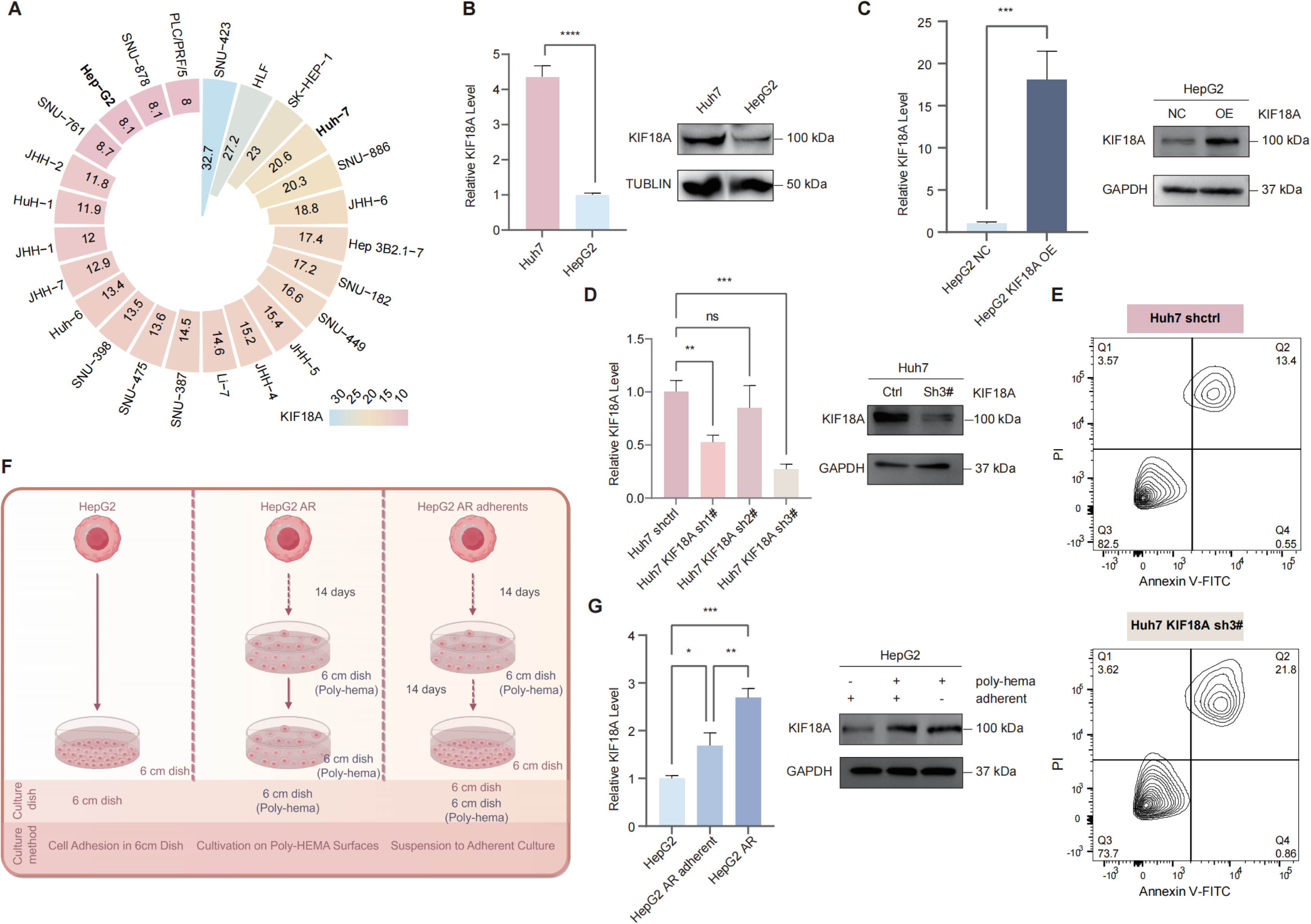
Investigation of KIF18A’s Role in Hepatocellular Carcinoma Cell Lines. (A) Basal expression levels of KIF18A in various hepatocellular carcinoma cell lines. (B) Immunoblotting and quantitative PCR analyses of KIF18A in HepG2 and Huh7 cell lines. (C) Relative KIF18A levels after overexpression in HepG2 cells with specific constructs. (D) Relative KIF18A levels after knockdown in Huh7 cells, and selected the most efficient knockdown of Huh7 KIF18A sh3#. (E) Apoptosis susceptibility in Huh7 KIF18A sh3# cells. (F) Acquisition process of HepG2 AR and stable HepG2 AR adherent cells. (G) KIF18A in HepG2 AR cells compared to HepG2 cells and HepG2 AR adherent cells.

To validate KIF18A’s role in anoikis resistance, we cultured HepG2 cells on ultra-low adhesion dishes utilizing poly-HEMA-coated dishes. After seven generations, we obtained HepG2 AR cells that exhibited resistance to anoikis and demonstrated stable survival. Furthermore, we cultivated HepG2 AR cells in standard adherent dishes for seven generations, resulting in stable HepG2 AR adherent cells (Fig. 7F). Results from transcription and protein level analyses indicated that KIF18A was upregulated in HepG2 AR cells relative to HepG2 cells, while re-adhered HepG2 AR cells continued to exhibit elevated KIF18A expression compared to HepG2 cells (Fig. 7G). This aligns with the findings of our single-cell research and the increase of KIF18A both prior to and following CTC conversion (Fig. 5B and 5C). HepG2 AR cells did not enter a dormant low KIF18A state, as observed in the PT-CTC and RT-CTC models. We hypothesize that the absence of complex cellular interaction and blood flow shear pressures in the culture is preventing the simulated conditions from attaining a fully inactive state for the CTCs. Therefore, further research is required to fully understand the CTCs behavior in the blood.

### The Multifaceted Role of KIF18A in Hepatocellular Carcinoma Metastasis and Its Association with the cGAS-STING Pathway

Previous studies on KIF18A have primarily focused on its role in WGD and CIN, which are key features of aneuploid tumors. Our research extends these findings by examining the ability of tumor cells to resist anoikis. We identified KIF18A as a hub gene associated with anoikis resistance, with many of its functions related to chromosomal dynamics (Fig. 1C, 3C, and 3F). In collaboration with Jan Lammerding’s research, we explore how to exacerbate the phenomenon of CIN^50^. Tumors with CIN tumors typically present a poorer prognosis and exhibit faster metastatic potential ^51^. We propose that anoikis resistance is an incremental consequence of CIN during tumor progression, with KIF18A acting as a key link between these two processes.

To further investigate this, we validated cGAS-STING activation in cell lines with overexpression (OE) and knockdown of KIF18A. HepG2 KIF18A OE cells showed elevated levels of p-STING and downstream p-TBK1 along with micronucleus formation (Fig. 8A and 8B). We hypothesize that increased micronuclei formation driven by

**Figure 8:**
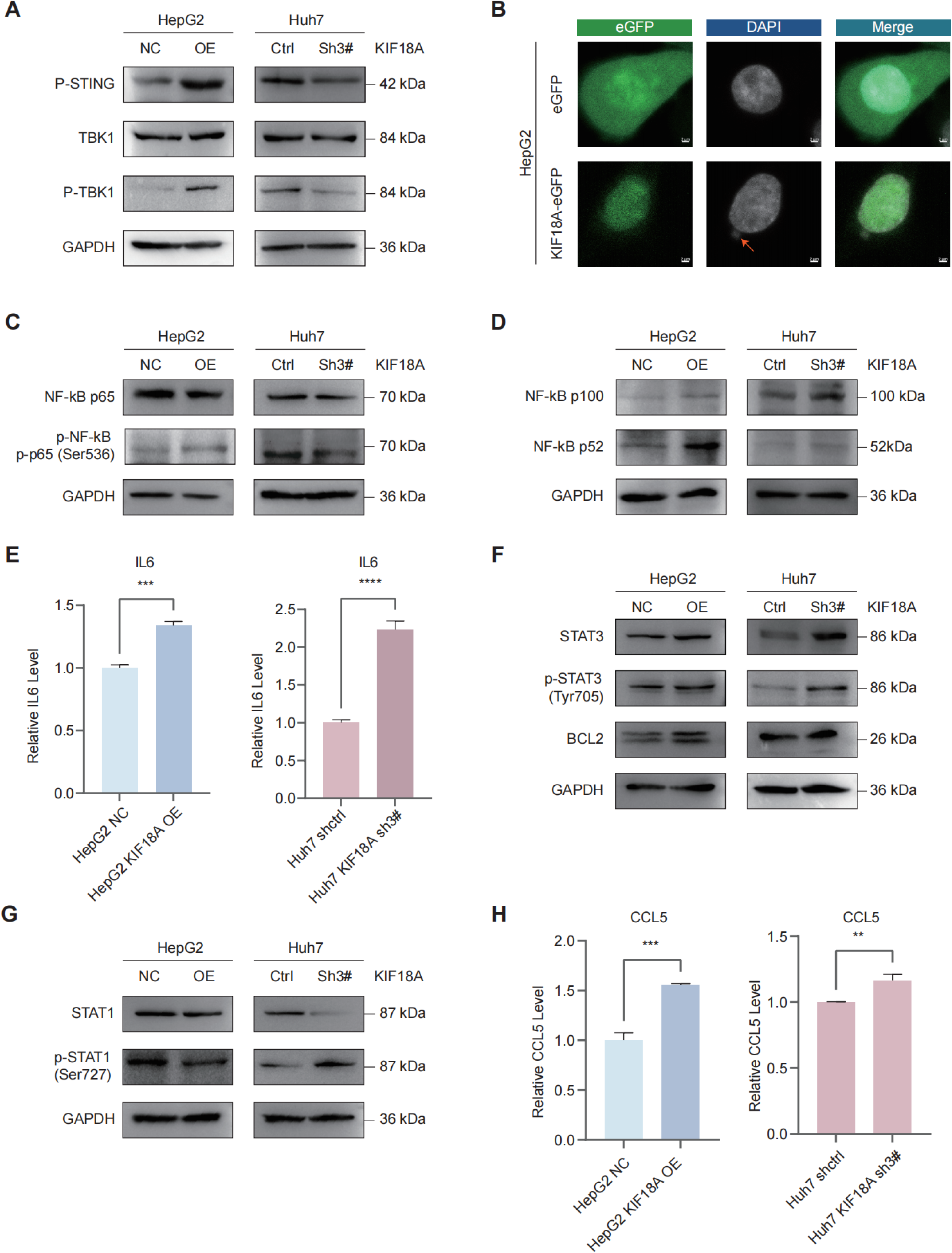
KIF18A association with the cGAS-STING Pathway. (A) Western blot analysis of cGAS-STING pathway activation in HepG2 KIF18A OE cells and Huh7 KIF18A sh3# cells. (B) Micronucleus formation in HepG2 KIF18A OE cells. (C) Western blot analysis of the canonical NF-kB pathway in HepG2 KIF18A OE cells and Huh7 KIF18A sh3# cells. (D) Western blot analysis of the non-canonical NF-kB pathway in HepG2 KIF18A OE cells and Huh7 KIF18A sh3# cells. (E) Quantitative analysis of IL6 levels in HepG2 KIF18A OE cells and Huh7 KIF18A sh3# cells. (F) Western blot analysis of STAT3 and BCL2 expression in HepG2 KIF18A OE cells and Huh7 KIF18A sh3# cells. (G) STAT1 activation in KIF18A-modulated cells. (H) Quantitative analysis of CCL5 levels in HepG2 KIF18A OE cells and Huh7 KIF18A sh3# cells.

WGD+/CIN fitness, combined with nuclear membrane rupture and the subsequent release of dsDNA, triggers chronic cGAS-STING activation^52^. Leslie A. Sepaniac et al. demonstrated that KIF18A ablation stabilizes the nuclear membrane, reducing the likelihood of nuclear membrane rupture and dsDNA release^53^. This finding aligns with our observation that knockdown of KIF18A inhibits cGAS-STING pathway activation in Huh7 KIF18A sh3# and Huh7 shctrl (Fig. 8A).

Moreover, when these four cell lines were cultured in poly-HEMA-coated dishes, HepG2 OE and Huh7 shctrl cells were more likely to form larger cultures of tumor islands compared to HepG2 NC and Huh7 KIF18A sh3# cells., which is a characteristic feature of tumor cells resistant to anoikis^54^ (Fig. S7C).

The cGAS-STING pathway is well-known for its anticancer effects in various cancers, promoting tumor metastasis when activated and inhibiting it when blocked ^55^. We first examined the canonical NF-kB signaling pathway downstream of cGAS-STING. In Huh7 KIF18A sh3# cells, p-NF-kB levels were downregulated due to the inhibition of STING phosphorylation, which led to the suppression of the canonical NF-kB pathway, which was significantly activated (Fig. 8C). Notably, in HepG2 KIF18A OE cells, the canonical NF-kB pathway was not triggered despite TBK1 phosphorylation (Fig. 8C); instead, the non-canonical NF-kB pathway was significantly activated (Fig. 8D). We propose that acute STING activation induces interferon release via classical pathway signaling, thereby enhancing antitumor immunity. In contrast, chronic STING activation shifts the response to nonclassical pathway activation, where non-canonical NF-kB (RELB-p52) is activated downstream of STING, antagonizing IRF3 and canonical NF-kB signaling while promoting the upregulation of genes associated with epithelial-mesenchymal transition and metastasis^56,57^.

IL6 is commonly considered a downstream marker of canonical NF-kB pathway activation^58^, and is widely believed to activate the IL6-JAK1-STAT3 signaling pathway to promote tumor survival. However, as shown in Fig. 8E and 8F, IL6-JAK1-STAT3 in HepG2 KIF18A OE cells is not activated, which is consistent with our single-cell SCPA data analysis results (Table. S4). Therefore, we hypothesize that while IL6-JAK1-STAT3 signaling pathway only ensures tumor cell survival^59^, it does not play a critical role in the formation of CTCs. Interestingly, in the Huh7 KIF18 sh3# cell line, which exhibited significantly increased apoptosis and inhibited canonical NF-kB signaling, IL6 mRNA levels as well as STAT3 and p-STAT3 protein levels were elevated but did not lead to BCL2 upregulation (Fig. 8F). Based on the research conclusions of Amy E. Schade et al.^60^, we believe that this may be due to the silencing of the PI3K-AKT pathway induced by KIF18Ash^61^, the activation of various components of the IL6-JAK1-STAT3 pathway, leading to the expression of BMF. Among these, BMF (a BH3-only protein) is the most significant pro-apoptotic gene and is induced in all drug-sensitive cell lines, ultimately promoting apoptosis^60^.

STAT1 is the primary mediator in response to IFN, with its downstream transcriptional response depending on the nature of the induced transcriptional complex, particularly the type and the duration of IFN exposure^62,63^. Acute type I interferon expression stimulates the production of interferon-stimulated genes that have cytotoxic and antiviral properties, but persistent interferon activation triggers the expression of IFN-related DNA damage resistance signatures (IRDs), which contribute to cancer cell resistance to DNA damage-based therapies^64^. As shown in Figure 8G in HepG2 KIF18A OE cells, although cGAS-STING is activated, the p-STAT1 response to type I interferon is reduced. We suggest that this is due to continuous STING activation, which enhances STAT1’s tendency to form the U-ISGF3 complex and promotes tumor progression^63^. In contrast, in Huh7 KIF18A sh3# cells, STAT1 levels were downregulated, but the apoptosis-associated p-STAT1 (Ser727) levels were elevated. This may result from the downregulation of IRDs transcription and increased apoptosis following the suppression of cGAS-STING. Additionally, we investigated the downstream CCL5 of cGAS-STING, which is linked to metastasis^14,65^ and corresponds with our single-cell analysis results (Fig. 6A-6E).

## 4. Discussion

We Initially confirmed the link between cell anoikis resistance and CIN in liver cancer using bulk RNA-seq (Fig. 1C and 1D). From this, we identified hub genes and selected KIF18A as a promising target for further investigation. Knowing that CTCs are resistant to anoikis^39^, we constructed a detailed model tracking the progression of liver cancer from its early stages to CTCs and subsequent recurrence. This model allowed us to identify a cell cluster resembling CTCs, which could potentially transform into actual CTC, providing insights into how tumors progress, whether by local growth or metastasis (Fig. 5B). Additionally, using a single-cell model, we found that CTC cells have the ability to mimic the phenotype of immune cells (Fig. 4A-4C). By integrating models of dissemination, dormancy, and colonization^39^, we confirmed the critical role of KIF18A activation in cancer spread (Fig. 5C). Furthermore, we identified two groups of cells pre-and post-metastasis, with elevated KIF18A levels that significantly activated the E2F signaling pathway (Fig. 5A and Table S4). This pathway is known to enhance the transcription of KIF18A^44,66^, hence reinforcing its essential role in metastasis, particularly regarding anoikis resistance at the single-cell level.

KIF18A, a kinesin motor protein, is crucial for spindle formation and has been linked to whole-genome doubling and chromosomal instability in aneuploid malignancies^36^. Our study supports the crucial role of KIF18A in HCC. The absence of KIF18A in liver cancer significantly enhances apoptosis and is crucial for anoikis resistance (Fig. 7E and S7C). KIF18A demonstrates tumor-specific selectivity, making it a viable target for therapeutic intervention in cancer^44,47^. Additionally, our findings provide new insights into CIN cancers and the cGAS-STING signaling pathway. KIF18A promotes tumor survival by chronically activating the cGAS-STING pathway and facilitating metastatic pathways such as EMT through non-canonical NF-kB (Fig. 8A and 8D). In contrast, KIF18A depletion activates the IL6-JAK1-STAT3 pathway but does not elevate BCL2, which is an anti-apoptotic protein (Figs. 8E and 8F). We hypothesize this is due to the inhibition of the PI3K-AKT pathway following KIF18A depletion^61^, aligning with the findings of Amy E. Schade et al^60^. STAT1 serves as the principal mediator in response to IFN, with its transcriptional response influenced by the type of IFN and exposure duration^62,63^. Acute IFN expression stimulates the production of cytotoxic and antiviral genes, while persistent activation triggers IFN-related DNA damage resistance signatures, associated with cancer cell resistance to DNA-damaging therapies^64^. In HepG2 KIF18A OE cells, although cGAS-STING is activated, the p-STAT1 response to type I interferon is reduced due to continuous STING activation, which promotes tumor progression, and in Huh7 KIF18A sh3# cells, STAT1 is downregulated, but p-STAT1 is elevated (Fig. 8G). This is likely attributable to the downregulation of IRDs transcription and heightened apoptosis^64^.

And as another downstream target of cGAS-STING, CCL5, which is closely related to metastasis^14,48^, is upregulated following KIF18A overexpression.

Our findings contribute to the construction of a liver cancer metastasis evolution model. In the early stages of liver cancer, some tumor cells exhibit abnormal activation of the E2F pathway, leading to increased transcription of KIF18A, which helps CIN tumors survive. This activation also chronically stimulates the cGAS-STING signaling pathway, promoting the non-canonical NF-kB rather than the canonical NF-kB pathway, thus controlling IFNγ production and promoting tumor metastasis. Additionally, apoptosis-related STAT1 was inhibited, IRDs-associated transcription was not activated, and CCL5 was partially upregulated.

Despite these significant findings, our study has some limitations. We only conducted cell experiments to explore the cGAS-STING and downstream pathways. More animal experiments are needed to validate our conclusions and fully understand the behavior of CTCs in the blood. Additionally, the downstream changes brought about by the transient activation of the cGAS-STING pathway require further investigation. We also need to determine whether the downstream pathways have other regulatory mechanisms and how the specific deletion of KI18A activates IL6-JAK1-STAT3 to induce BMF-related apoptosis.

This study examined the relationship between anoikis resistance and CIN in HCC using bulk RNA sequencing and single-cell RNA sequencing technologies. We identified the KIF18A gene as a pivotal hub affecting these two essential pathways. Our research indicates that KIF18A may promote tumor metastasis by stimulating the cGAS-STING signaling pathway, thus affecting the expression of CCL5. KIF18A may facilitate tumor cell spread by assisting circulating tumor cells in platelet recruitment via CCL5, so increasing their resilience to shear stress in blood flow and strengthening their resistance to anoikis and DNA damage^67^(Fig. 9). These discoveries emphasize the potential of KIF18A as a target for novel therapeutic strategies against HCC and underscore its critical role in the progression and metastasis of HCC. Future study will focus on clarifying the specific mechanisms of KIF18A function and investigating methods to effectively target it to reduce metastasis and improve patient outcomes.

**Summary Figure (Fig. 9).**
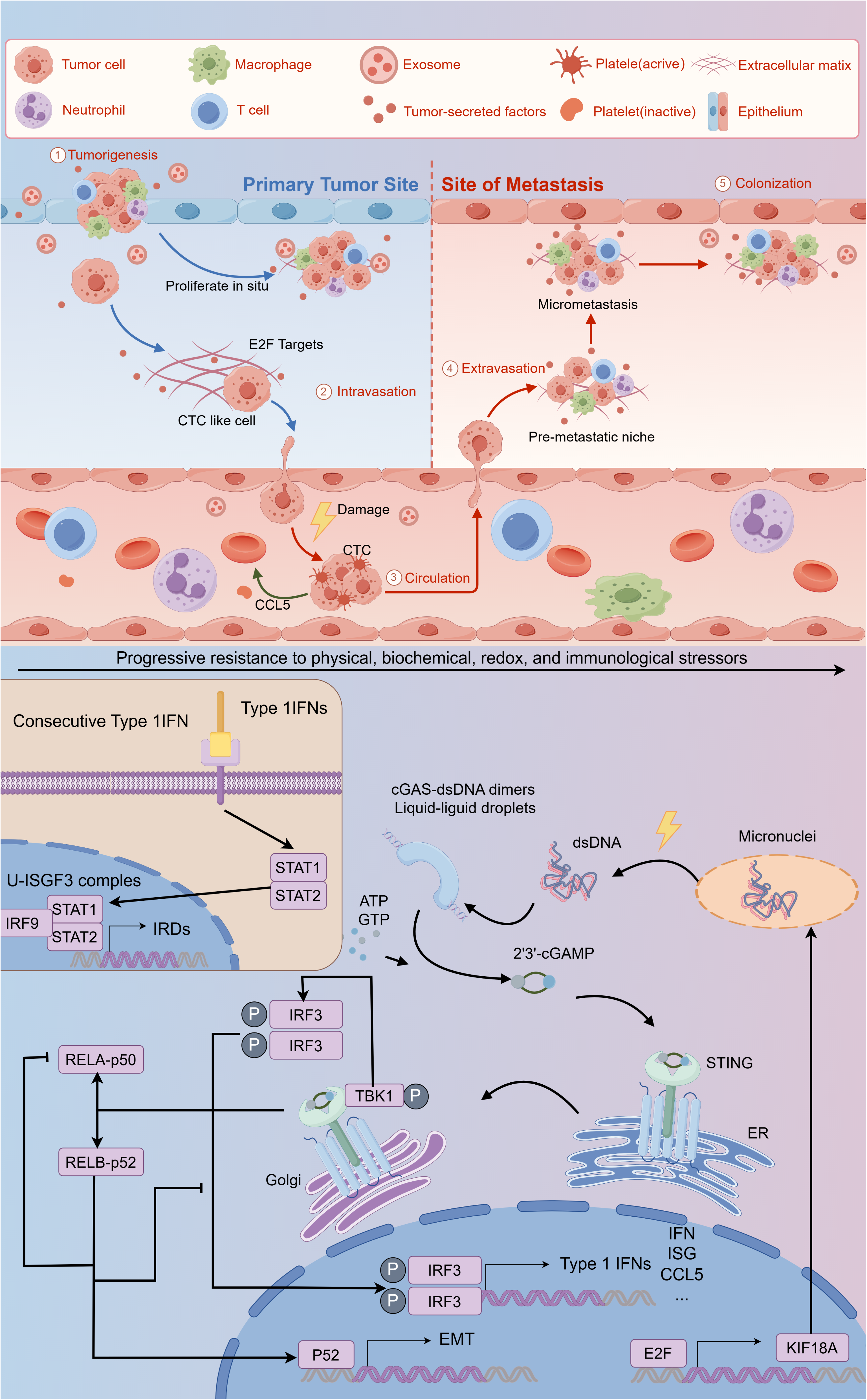

## Data availability statement

Publicly available datasets were analyzed in this study. The datasets used and analyzed in the study are available from the corresponding author on reasonable request.

## Acknowledgment

We thank Dr. Jing Li for their helpful suggestion and comments on the manuscript. We thank the Center for Life Sciences, Yunan University for providing the experimental platform. We thank Youzhou Li (Flow Cytometry Platform, Life Science Center, Yunnan University) for his help. We thank Dr.Jianming Zeng(University of Macau), and all the members of his bioinformatics team, biotrainee, for generously sharing their experience and codes. The Use of the biorstudio high performance computing cluster(https://biorstudio.cloud) at Biotrainee and The shanghai HS Biotech Co.,Ltd for conducting the research reported in this paper. Since Biomamba and his wechat public account team produce bioinformatics tutorials elaborately and share code with annotation, we thank Biomamba for their guidance in bioinformatics and data analysis for the current study.

## Supplementary Figures

**Figure S1: Cluster Analysis and Survival Probability of ARGs in HCC**

(A) Venn diagram displaying the overlap of ARGs between differentially expressed genes in the TCGA-LIHC dataset and those listed in the GeneCards database. (B) Consensus clustering analysis within HCC samples to determine cluster optimality. (C) Consensus Index Plot for identifying the optimal number of clusters in HCC samples. (D) PCA plot showing the separation of samples into two distinct subtypes, with Cluster 1 represented by red spots and Cluster 2 by blue spots. (E) Kaplan-Meier survival curves for the two clusters (C1 and C2), illustrating survival probabilities over time. (F) Distribution of patient status and tumor stages across different clusters, as indicated by a summary table or plot.

**Figure S2: Cellular Proportions and Pathway Analysis in Groups C1 and C2**

(A) Relative proportions of dendritic cells, lymphocytes, macrophages, and mast cells in groups C1 and C2, with statistically significant differences highlighted. (B) Comparison of tumor microenvironment scores, including Stromal Score, Immune Score, ESTIMATE Score, and Tumor Purity, between groups C1 and C2. (C) Proportions of various immune cell subsets, such as naive B cells, memory B cells, plasma cells, and CD8+ T cells, in groups C1 and C2. (D) GSVA scores comparing the activity of Hallmark pathways between groups C1 and C2, showing significant differences in pathway activity.

**Figure S3: Gene Identification and Decision Curve Analysis for LIHC Prognosis**

(A) LASSO regression analysis identifying six genes (KIF18A, SPP1, PLK1, SLC2A2, EZH2, MMP3) significantly associated with survival outcomes in patients with LIHC. (B) Relationship between patient status and gene expression, illustrating the association between patient status, clusters of ARGs, and ARS. (C) Increased ARS observed in patients with the ARPGs-C1 subtype. (D) Decision Curve Analysis (DCA) for the Nomogram over 1, 3, and 5 years, assessing the clinical utility of the prognostic model.

**Figure S4: Cellular Analysis in HCC Samples**

(A) UMAP plot of HCC cells, including adjacent liver, CTC, and tumor cells, providing a comprehensive view of cellular heterogeneity. (B) Proportional distribution of cell types within HCC samples, including T cells, macrophages, hepatocytes, CTC-like cells, B cells, and endothelial cells. (C) Manhattan plot of gene expression for the top 5 upregulated and downregulated genes in each cluster. (D) inferCNV analysis of CTC-like primary cells for CNVs, offering insights into the genomic alterations associated with these cells.

**Figure S5: Reclassification and Trajectory Analysis of CTC-like and Liver Cells in HCC**

(A) UMAP plot of PT-CTC and RT-CTC, revisiting the clustering of hepatocyte and CTC-like cells from the previous analysis. (B) UMAP plot identifying the area of cells within the PT-CTC and RT-CTC models, distinguishing cells derived from tumor, CTC, and adjacent liver tissues. (C) Trajectory analysis using Slingshot and k-means clustering, showing the differentiation paths of cells, with CTC serving as either the endpoint in PT-CTC or the starting point in RT-CTC.

**Figure S6: Analysis of CCL Signaling Pathway in HCC Models**

(A)Outgoing and Incoming Signaling Patterns involving various mediators such as MK, PARs, MIF, ANGPTL, and others in PT-CTC of HCC models. (B) Cellular Roles in the CCL Signaling Pathway based on their roles in the CCL signaling pathway. (C) Cellular Roles in the CCL Signaling Pathway based on their roles in the CCL signaling pathway, this panel distinguishes between senders, receivers, and influencers.

**Figure S7: Investigation of KIF18A’s Role in Hepatocellular Carcinoma Cell Lines**

(A) Relative KIF18A protein levels in HepG2 cells following transfection with different shRNA constructs targeting KIF18A.(B) KIF18A overexpression in Huh7 cells. (C) Tumor island formation analysis, comparing the cultivation of four cell lines in poly-HEMA-coated dishes, including HepG2 overexpressing KIF18A (OE) and Huh7 cells with control shRNA (shctrl).

