## Supplementary figures and images for "KIF18A in HCC Metastasis: Dual Role in Anoikis Resistance and Chromosome Instability"

### Supplemental Figure 1

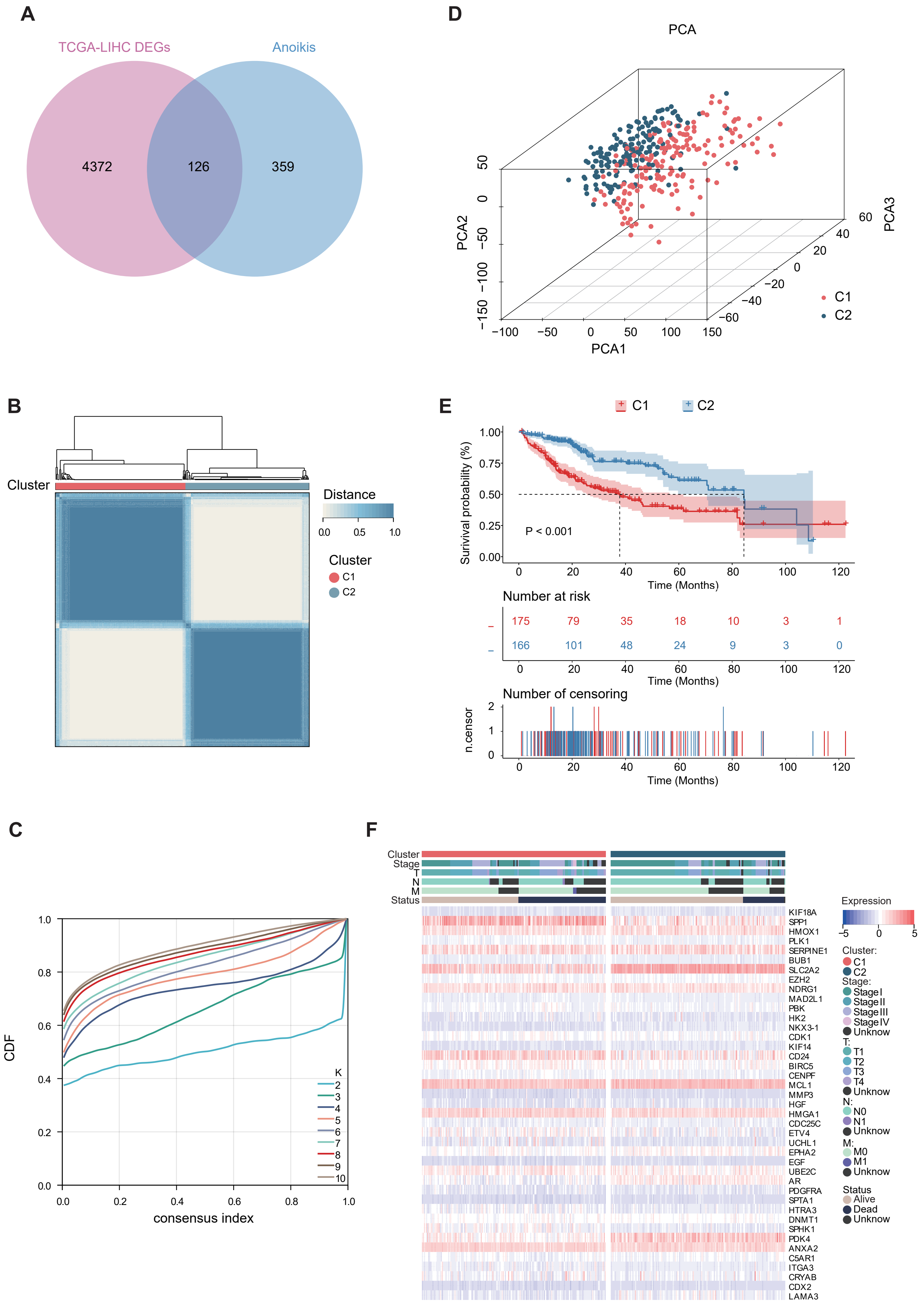

### Supplemental Figure 2

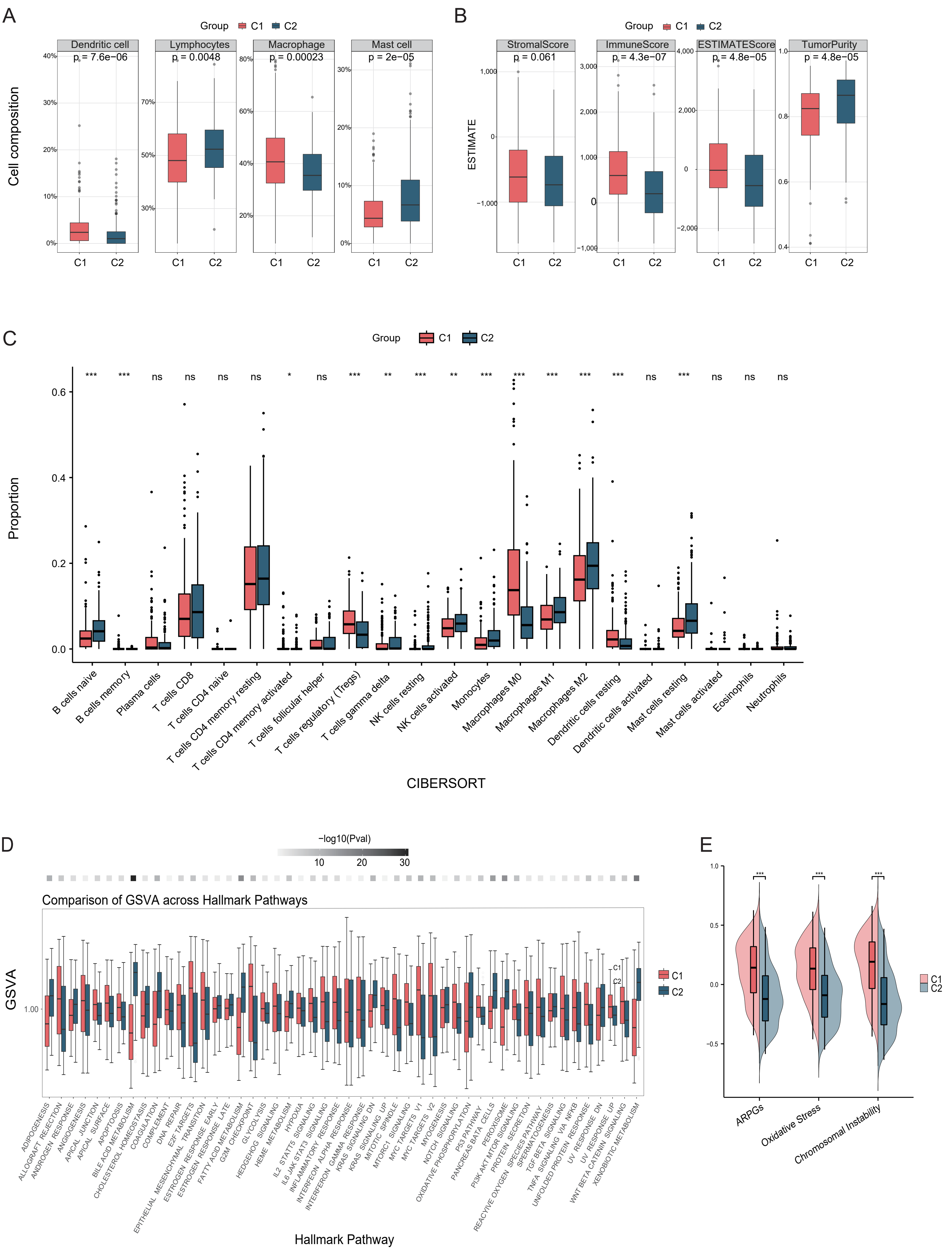

### Supplemental Figure 3

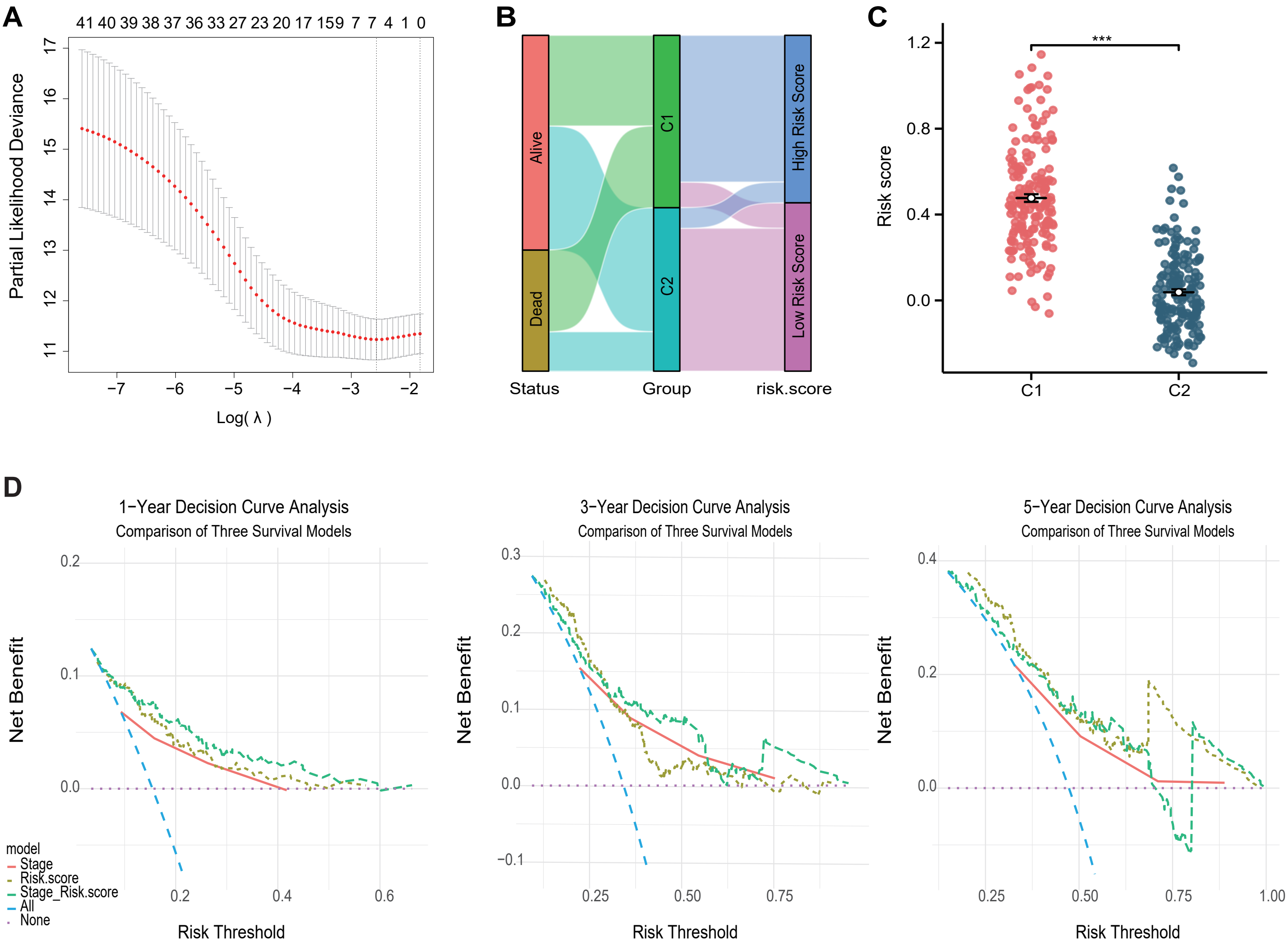

### Supplemental Figure 4

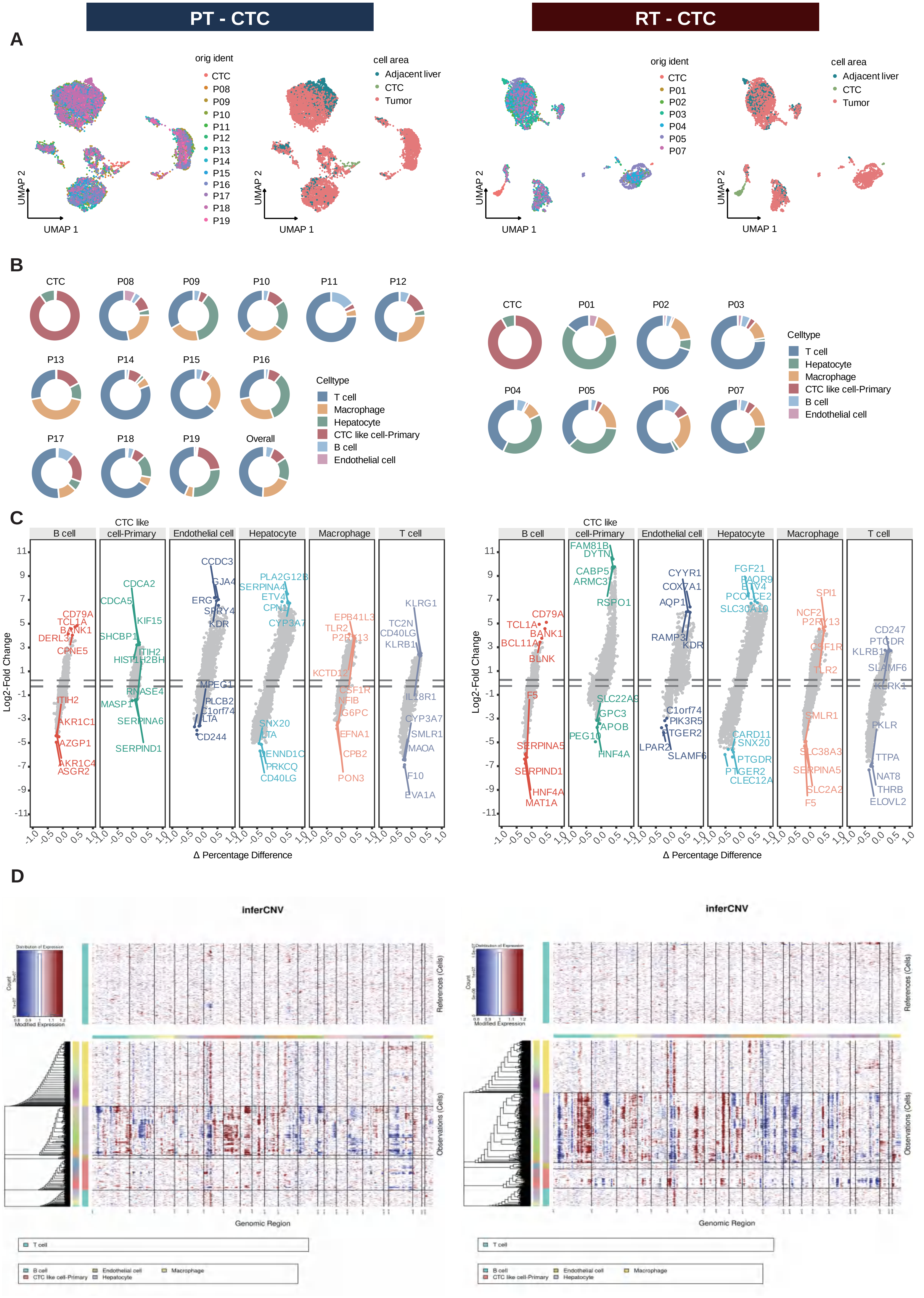

### Supplemental Figure 5

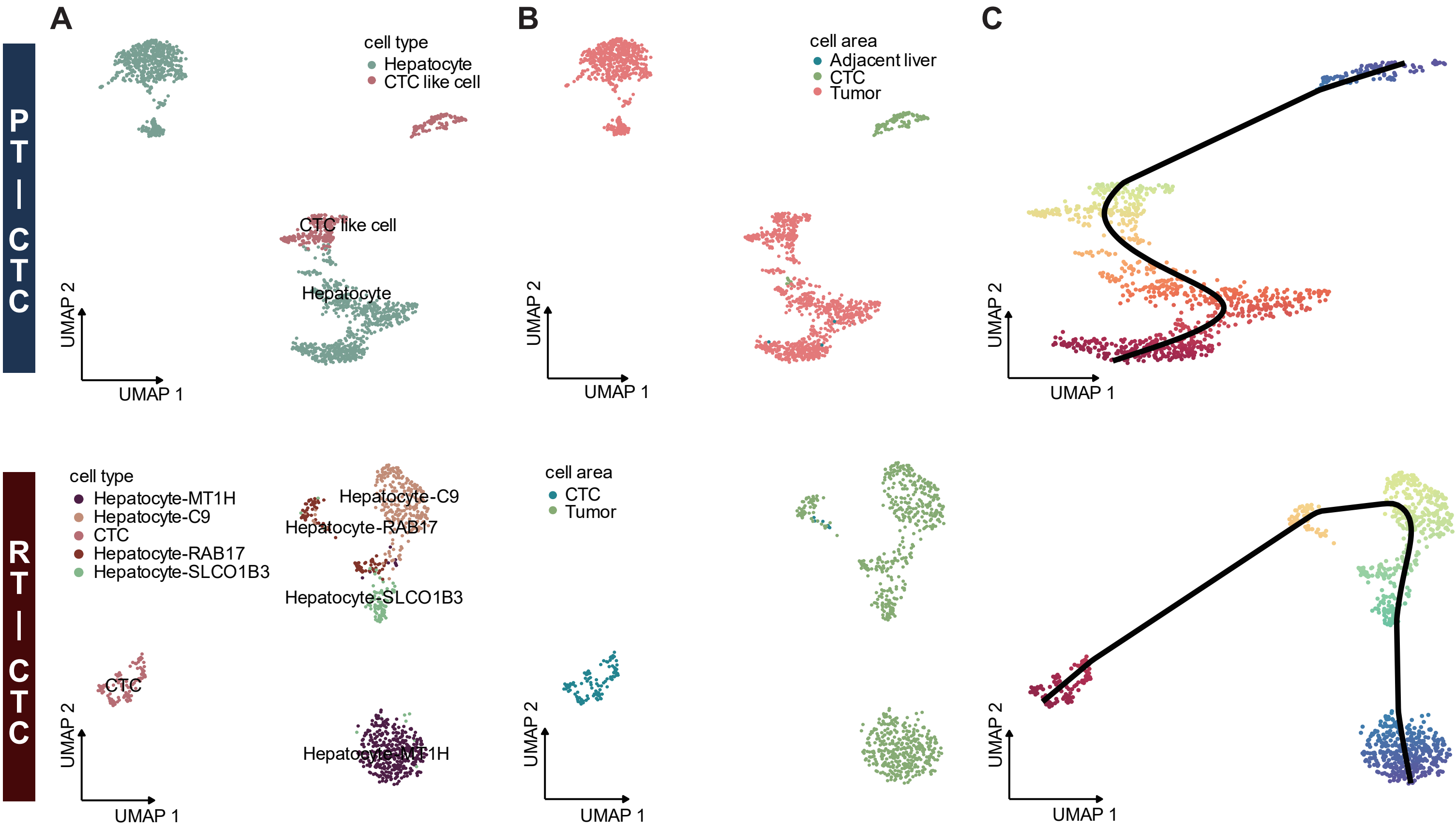

### Supplemental Figure 6

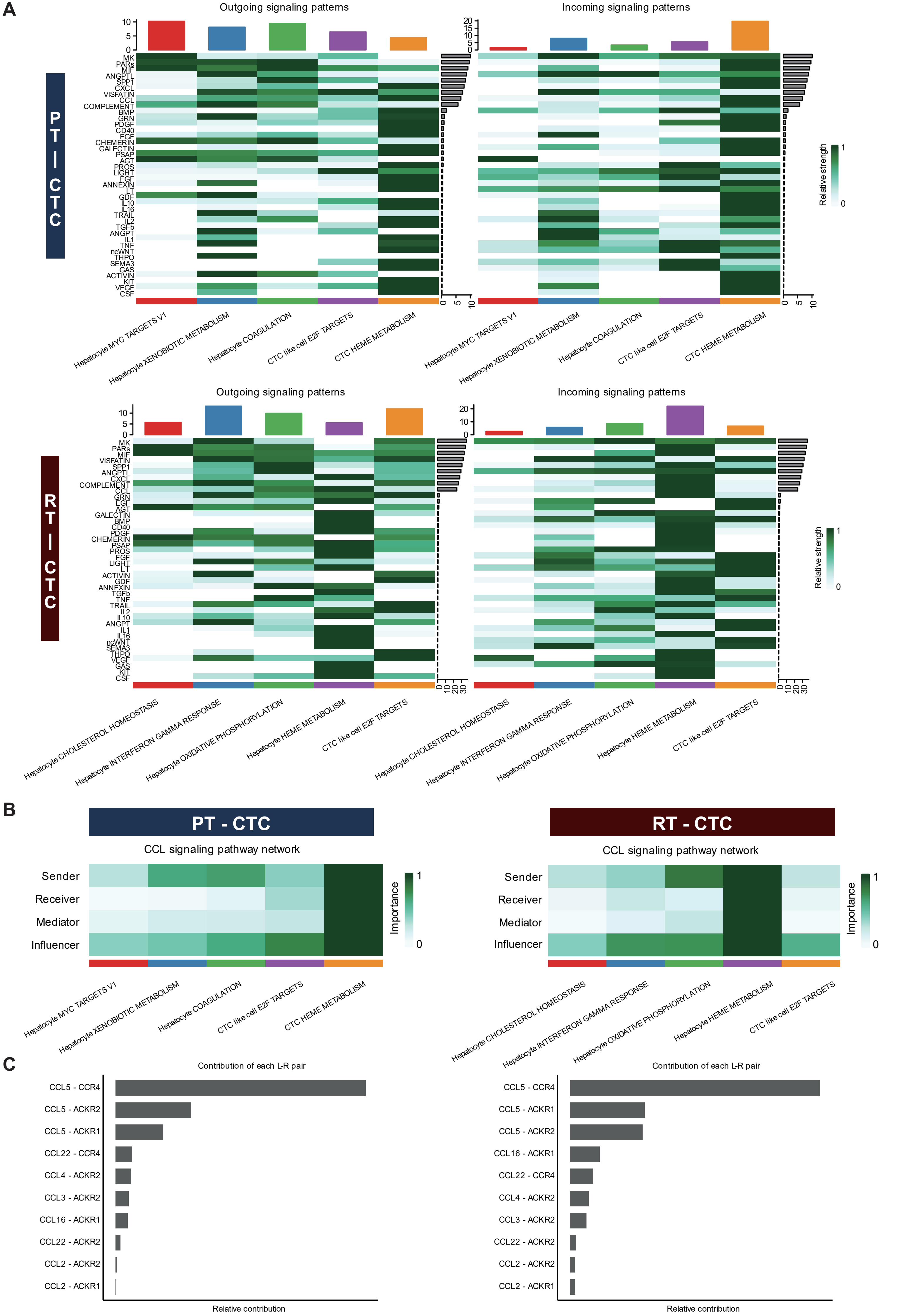

### Supplemental Figure 7

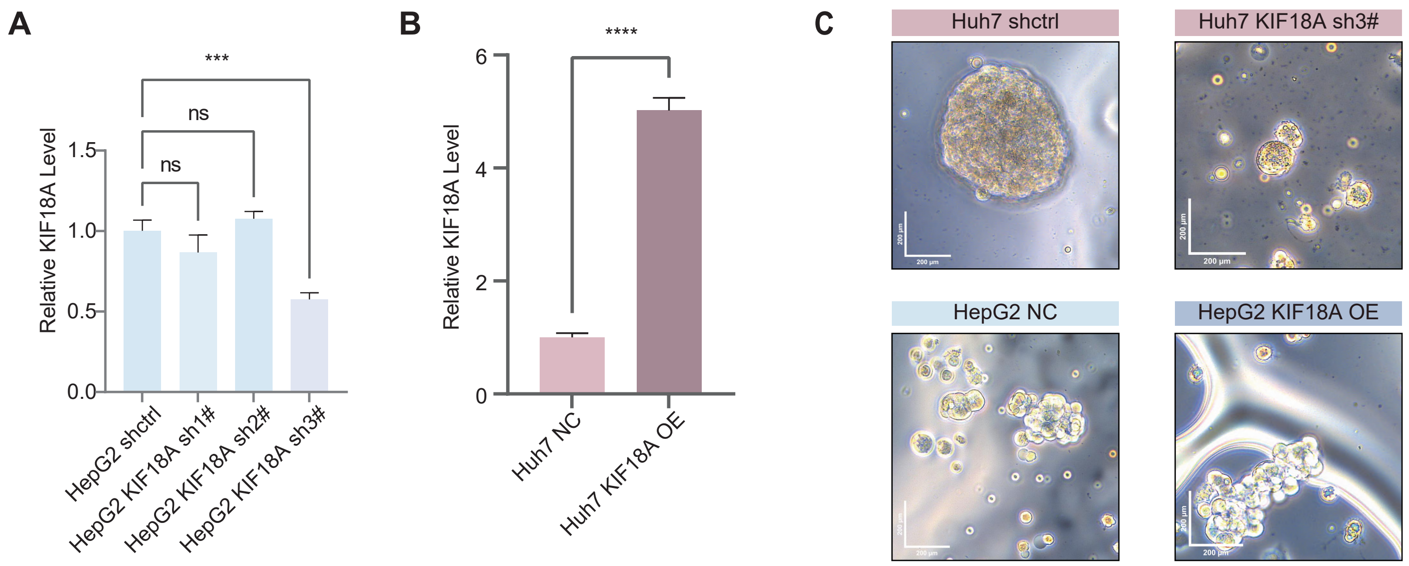
